# Unraveling nanotopography of cell surface receptors

**DOI:** 10.1101/2019.12.23.884460

**Authors:** Christian Franke, Tomáš Chum, Zuzana Kvíčalová, Daniela Glatzová, Alvaro Rodriguez, Dominic A. Helmerich, Otakar Frank, Tomáš Brdička, Sebastian van de Linde, Marek Cebecauer

## Abstract

Cells communicate with their environment via surface receptors, but nanoscopic receptor organization with respect to complex cell surface morphology remains unclear. This is mainly due to a lack of accessible, robust and high-resolution methods. Here, we present an approach for mapping the topography of receptors at the cell surface with nanometer precision. The method involves coating glass coverslips with glycine, which preserves the fine membrane morphology while allowing immobilized cells to be positioned close to the optical surface. We developed an advanced and simplified algorithm for the analysis of single-molecule localization data acquired in a biplane detection scheme. These advancements enable direct and quantitative mapping of protein distribution on ruffled plasma membranes with near isotropic 3D nanometer resolution. As demonstrated successfully for CD4 and CD45 receptors, the described workflow is a straightforward quantitative technique to study molecules and their interactions at the complex surface nanomorphology of differentiated metazoan cells.

## INTRODUCTION

Supramolecular complexes drive numerous vital processes in cells, such as gene expression, molecular transport or signal transduction. Cellular membranes provide an excellent platform to assemble molecules into complex structures. Indeed, membrane-associated molecules, including surface receptors, have been found to form clusters with nanometric dimensions^1–5^. Lateral interactions and/or actin-cytoskeleton anchorage drive clustering of some receptors (e.g., tetraspanin platforms, focal adhesion complexes). However, the mechanism of cluster assembly for other receptors remains unknown and proposed theories are often controversial (e.g., receptor microclusters on lymphocytes). Indeed, receptor clustering is the subject of intense debate^6–8^. Several works have provided evidence for the monomeric character of receptors, which were shown to cluster using different experimental setups^9–11^. Thus, understanding their origins can help to unravel the very existence of receptor clusters. In this work, we focused on receptor clustering on the plasma membrane of lymphocytes, but the method and general principles discussed herein apply to the surface of any cell, including those of prokaryotes.

A common feature of supramolecular structures, including membrane receptor clusters, is their size (< 200 nm), too small to be analyzed by standard light microscopy. Therefore, molecular assemblies in cells are often studied indirectly. For example, total internal reflection fluorescence (TIRF) microscopy enabled dynamic receptor microclusters to be visualized in B and T cells^12,13^. It was the lateral mobility of these entities that indicated the very existence of clusters in these cells. The size and shape of the observed clusters were irresolvable because the spatial resolution of TIRF microscopy is diffraction limited. Later, super-resolution (SR) microscopy techniques were developed to surpass the diffraction limit and offered a more detailed insight into the architecture of cell receptor clusters (for example, refs: ^5,14–16^).

Currently, single-molecule localization microscopy (SMLM) is the method of choice to study the organization of membrane receptors due to its ability to localize emitters (i.e., labelled receptors) with nanometer precision and its potential for quantitative assessments of protein distribution and number^17–20^. SMLM-based studies confirmed the earlier observations using TIRF microscopy that receptors and associated signaling molecules can cluster in non-stimulated and stimulated immune cells^21–25^. However, there is an intense discussion about the feasibility of SMLM for the cluster analysis of membrane molecules^26–28^. Recently, procedures aiming to minimize the impact of methodological artefacts on the SMLM results were developed^29–31^.

In most of these studies, the SMLM methods were used to generate localization maps by projecting the presumably three-dimensional receptor distribution onto a two-dimensional plane. The precise information about the axial position of emitters (e.g., receptors), and therefore their distribution, is missing. This was an accepted trade-off due to the limited axial resolution of the applied SR methods (including STED and SIM; ref.^32^). Moreover, the plasma membrane is routinely depicted as a rather featureless structure (for example, refs.^33,34^). Yet differentiated cells of vertebrates are densely covered with membrane protrusions and invaginations, thus resulting in a highly three-dimensional surface^35^. For example, scanning electron microscopy micrographs have demonstrated that finger-like membrane protrusions reminiscent of microvilli dominate the surface of lymphocytes^36–38^. Microvilli are dynamic structures ~100 nm in diameter and 0.5–5 μm long^39^. Although less well understood on immune cells, tips of microvilli can potentially accumulate membrane receptors in domains with a diameter ~100 nm, analogous to signaling receptors and channels on epithelial and sensory cells^40^.

Electron microscopy (EM), with its ability to provide information about ultrastructural details, was the key method for microvilli characterization in the pioneering works^41^. However, EM cannot visualize specific proteins efficiently, since the labelling densities are limited by ligand/antigen accessibility, steric hindrance and electron repulsion^42–44^. A rapid development of SR light microscopy techniques enabled the visualization of three-dimensional objects with high precision, with the use of highly specific and frequently efficient labelling methods and on living cells (e.g., refs.^45,46^). However, to characterize receptor distribution on nanometric membrane structures such as microvilli, it is crucial to develop three-dimensional SR techniques with axial resolution well below 50 nm. Several SMLM based methods have been suggested^47^, such as astigmatism^48^, multiplane^46^ or biplane imaging^49,50^, double-helix point spread function^51^ or interferometric PALM^52^.

Although microscopy techniques with dramatically improved spatial resolution recently became available, the (cell) sample preparation for SR imaging still suffers from several caveats. The spatial resolution offered by SMLM comes with the tradeoff of time resolution since thousands of camera frames are needed to render a map sufficiently representing, for example, a receptor distribution on a cell^53^. Thus, the movement of imaged objects (e.g., proteins) has to be minimized. This is usually accomplished by the fixation of cells prior to imaging. Importantly, cells grown or isolated initially in suspension must be attached to the optical surface (e.g., coverslips). Poly-*L*-lysine (PLL, or its isomer poly-*D*-lysine) is used for the immobilization of suspension cells in standard protocols^54^. However, attachment to a surface via interaction with PLL leads to the deformation of cells^55^. Similarly, the flattening of cell surfaces is observed when positioning immune cells on adhesion molecules (e.g., ICAMs) directly coated on coverslips or linked to a supported planar lipid bilayer^56,57^. The use of hydrogel or Matrigel enables cell stabilization for imaging without observable morphological changes^58,59^. However, these materials submerge cells into a three-dimensional matrix which leads to their random axial distribution. Single-molecule fluorescence microscopy techniques (including SMLM) depend on efficient transmission of photons from emitters and thus, provide the best results for molecules close to the optical surface, especially when aiming for a subsequent quantitative analysis^47^. Currently, none of the available methods can immobilize cells close to the optical surface without interfering with their complex surface morphology.

CD4 is a surface glycoprotein involved in T-cell development and function. It is associated with Lck kinase, which drives antigen-specific T-cell activation^60,61^. CD4 also mediates T-cell adhesion to the cells which express major histocompatibility complex class II (MHCII) proteins on their surface. We have recently shown that CD4 accumulates in clusters on the surface of non-stimulated T cells in a palmitoylation-dependent manner^28^. The origin of these CD4 clusters remains unknown. CD45, another glycoprotein that is three times larger than CD4, is expressed at the surface of all lymphocytes^62^. Its intracellular phosphatase domain regulates Lck kinase activity. CD45 has been found to segregate from signaling molecules in activated T cells, as predicted by the kinetic segregation model of T-cell activation^63,64^. However, the mechanism responsible for such segregation is incompletely understood due to a lack of information about its precise localization at the surface of immune cells.

Here, we describe a new method for mapping receptors at the nano-topography of the plasma membrane with near isotropic three-dimensional resolution. The workflow involves a newly developed, optimized coating of glass coverslips, which preserves the complex morphology of a cell surface while allowing for the positioning of immobilized cells close to the optical surface. Moreover, we advanced the recently reported TRABI method^65^ for nanometer precise three-dimensional SMLM imaging to improve data acquisition, processing and drift correction, as well as the integrity of the three-dimensional SMLM data set. This allowed us to employ a straightforward quantitative assessment of the axial distributions of selected cell surface receptors at the nanoscale. We applied this method to reveal the three-dimensional nature of CD4 receptor clusters in non-stimulated T cells. Also, we discovered the importance of membrane protrusions for the segregation of CD45 phosphatase from CD4 involved in T-cell signaling. The presented method enables molecular studies of cell surface nano-morphology but also of other fluorescent nanoscopic structures.

## RESULTS

### Preservation of membrane morphology and resting state during cell immobilization on coverslips

To map receptor distribution at the complex surface of cells and the nanoscale, the experimental setup must include quantitative labelling of target molecules, an appropriate sample preparation, and a nanometer precise three-dimensional imaging method. In this work, labeling of surface receptors was performed by standard immunofluorescence protocols using directly labelled, highly specific primary antibodies to avoid artificial clustering by high level crosslinking with secondary antibodies^66–68^. To prevent visualization of intracellular molecules, fixed T cells were labelled in the absence of membrane permeabilization.

Cells grown or isolated in suspension, e.g., T cells, must be immobilized on the optical surface (e.g., coverslips; Fig. 1a-b) to facilitate imaging approaches, which require long acquisition times. This is true for most SMLM techniques^69^. Commonly, PLL-coating of glass coverslips is used to adhere suspension cells to the optical surface^55^. Negatively charged biomolecules at the cell surface electrostatically interact with the polycationic layer formed by PLL on coverslips^70,71^. However, such interaction can deform the cell surface (Fig. 1a; ref.^55^). Indeed, we have repeatedly observed a rapid flattening of a T-cell surface upon settling on the PLL-coated coverslip, visualized by TIRF microscopy of CD4-GFP fusion protein (for example, Fig. 1c and Supplementary Movie 1). With small exceptions (red arrowhead), a homogenous distribution of CD4-GFP signal dominated in such cells within 1-5 minutes after the first detectable contact with the optical surface (blue arrowheads; and Supplementary Fig. S1a). Continuous and homogenous fluorescence signal indicated a lack of CD4-GFP accumulation on membrane protrusions and invaginations. These data suggest that immobilization on PLL-coated coverslips can cause the flattening of the T-cell surface.

**Figure 1.**
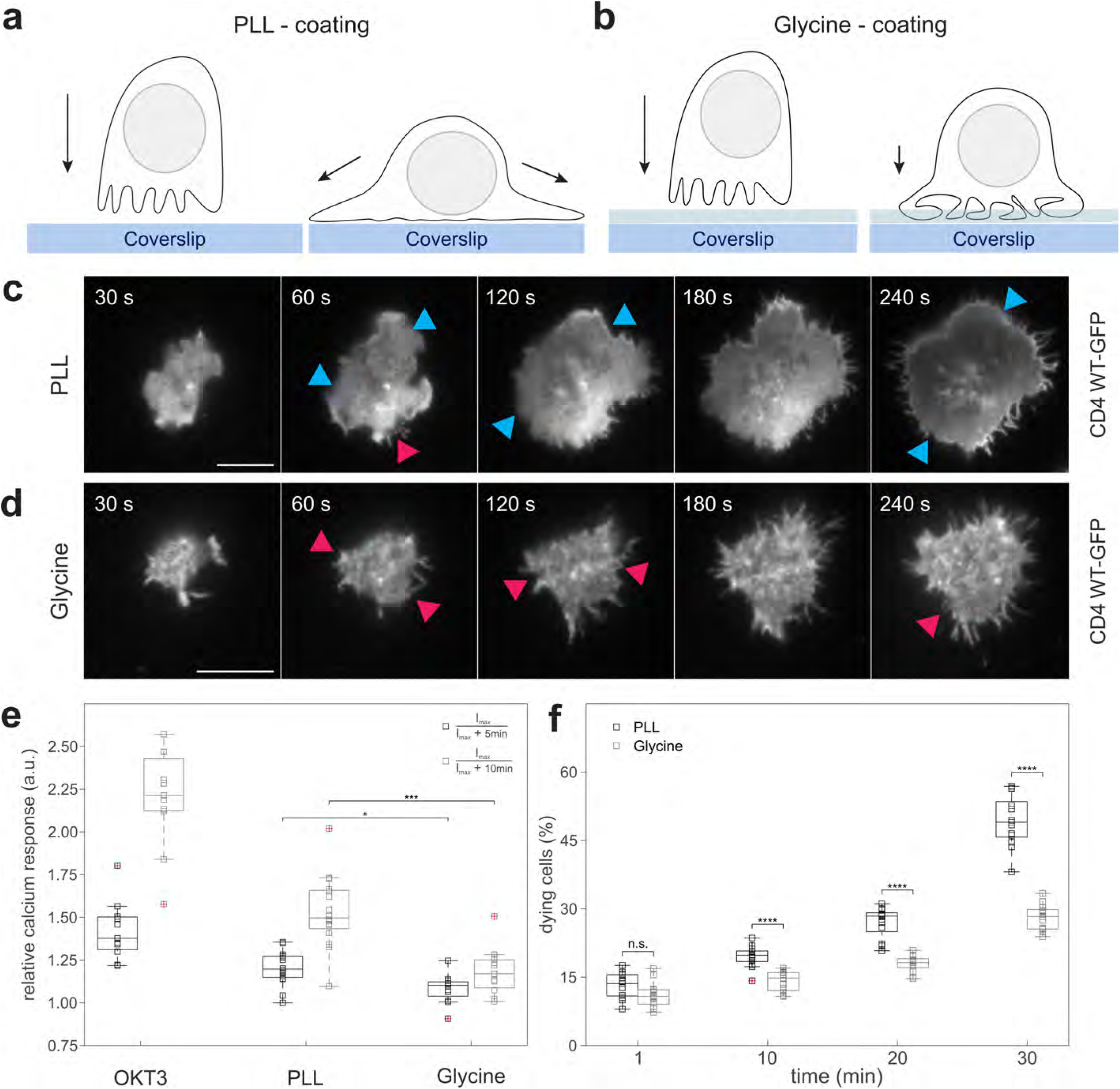
Coating of coverslips with glycine better preserves cell surface morphology and resting state of immobilized T cells. **a-b**) Schematic illustration of T cell landing on PLL-(left panels) and glycine-coated coverslips (right panels). The arrows indicate forces influencing the cell on the coated coverslip. The g-force is the sole force affecting T cells on glycine. T cells on PLL are further stretched due to electrostatic interactions of the surface molecules with PLL. The blue stripes represent glass coverslip, the light blue stripes above represent the glycine layer (not to scale). **c-d)** Live-cell TIRF microscopy of CD4-GFP in T cells landing on PLL-coated (c) or glycine-coated coverslips (d) measured at 37°C (see also Supplementary Movies 1 and 2). Blue arrowheads indicate areas of rapid flattening of the cell surface and random distribution of CD4, as indicated by line-profiles in Supplementary Fig. S1. Red arrowheads indicate areas with heterogeneous distribution of CD4 on the cell surface as indicated by line-profiles in Supplementary Fig. S1. Selected time points for the representative cells are shown. In total, 14 cells on glycine-coated and 10 cells on PLL-coated coverslips were analyzed. **e)** Calcium response induced by the interaction of Jurkat T cells with coverslips coated with stimulating antibody (anti-CD3ε; OKT3), PLL or glycine as indicated by the changes in ultrafast genetically encoded calcium sensor GCaMP6fu fluorescence over 5 min (black bars) and 10 min (grey bars) from the maximal stimulation of cells. Relative calcium response equal 1 indicates no stimulation. Higher values indicate stimulation of cells (see **Methods** and Supplementary Fig. S3 for more details). **f)** Viability of Jurkat T cells interacting with coverslips coated with PLL-(black bars) or glycine-coated (grey bars) coverslips. Dying cells were defined as a fraction of 7-aminoactinomycin-positive cells within the imaged area using wide-field microscopy (see **Methods** for more details). Values at 1 min represent the starting point for the analysis – a minimal period required for cells to land at the optical surface.

We thus required an alternative method for coating of coverslips, which better preserves their native morphology. Coating of an optical surface with glycine has been previously used to minimize non-specific background signals in single-molecule fluorescence imaging^72,73^. However, when applied without prior PLL treatment, glycine formed a gel-like layer on a glass surface (Supplementary Fig. S2a). Our atomic force microscopy (AFM) measurements indicated a continuous surface coating with glycine (Supplementary Fig. S2b). However, the AFM measurements were limited by the softness of the glycine layer and a tendency of this material to adhere to the AFM tip. We were thus unable to determine the exact thickness and stiffness of the glycine layer. The few holes, probably caused by gas bubbles, observed in the tested samples indicated that the average thickness of the glycine layer was at least 15 nm (Supplementary Fig. S2c). We further observed crystals on coverslips coated with glycine that were subsequently dried. On the contrary, no such precipitate was observed on dried PLL-coated coverslips (Supplementary Fig. S2d). We thus conclude that glycine forms a narrow, gel-like structure on the glass surface, which functions as a semi-soft cushion upon cell landing.

To investigate the impact of glycine coating on cell surface morphology, we imaged T cells expressing CD4-GFP during landing on coverslips in analogy to the cells on PLL-coated coverslips (Fig. 1c). Interestingly, a highly heterogeneous CD4-GFP signal was detected in a majority of cells immobilized on glycine-coated coverslips for more than 5 minutes since the first contact with the optical surface (Fig. 1d and Supplementary Movie 2). Intensity line profiles measured across the surface of such T cells exhibited large deviations (Supplementary Fig. S1b). These data indicate that glycine coating allows immobilization of cells close to the optical surface and improves the preservation of cellular morphology better, and for longer, than PLL-coating. Such properties open access to live-cell imaging of cell surface morphology or more convenient sample preparation, e.g., fixation.

We further examined the impact of the newly developed coating method on the resting state and viability of immobilized T cells. PLL was reported to induce calcium response in cells attached to the coated surface^74^. Our measurements confirmed that PLL induces calcium mobilization in T cells, albeit much less than antigenic stimulation mimicked by the coverslip coated with specific antibodies (anti-CD3ε) (Fig. 1e and Supplementary Fig. S3). In turn, the immobilization of T cells on glycine-coated coverslips stimulated only a negligible calcium response (Fig. 1e and Supplementary Fig. S3). We have employed the highly-sensitive, genetically encoded membrane-associated fluorescent calcium indicator GCaMP6f_u_, which enables detection of weak and rapid signals from cell surface receptors and channels^75^. Glycine-coating of coverslips, thus, prevents non-specific stimulation of T cells observed on PLL-coated surfaces. Moreover, for longer cultivation times (> 10 min) we found a significantly improved viability for T cells incubated on glycine-coated coverslips compared to those on PLL-coated coverslips (Fig. 1f).

In summary, our findings indicate that coating of optical surfaces with glycine reduces the stress generated by the charged surface of, for example, PLL-coated coverslips. Moreover, the gel-like structure of the glycine coating better preserves the surface morphology of immobilized cells and enables quantitative analysis of the three-dimensional receptor distribution on the surface of resting T cells with SMLM.

### The three-dimensional SMLM method

To capture the entire nanoscopic three-dimensional organization of receptors at the cell surface, we acquired image sequences with a highly inclined and laminated optical sheet (HILO) illumination, which enables single-molecule detection in cells much deeper than in TIRF microscopy (Fig. 2a). At the same time, it restricts the out-of-focus fluorescence from the remaining parts of a cell, thus improving the local signal-to-noise ratio^76^. Prior to the acquisition, we briefly irradiated the whole cell in epifluorescence to further minimize out-of-focus contributions (see **Methods**).

**Figure 2.**
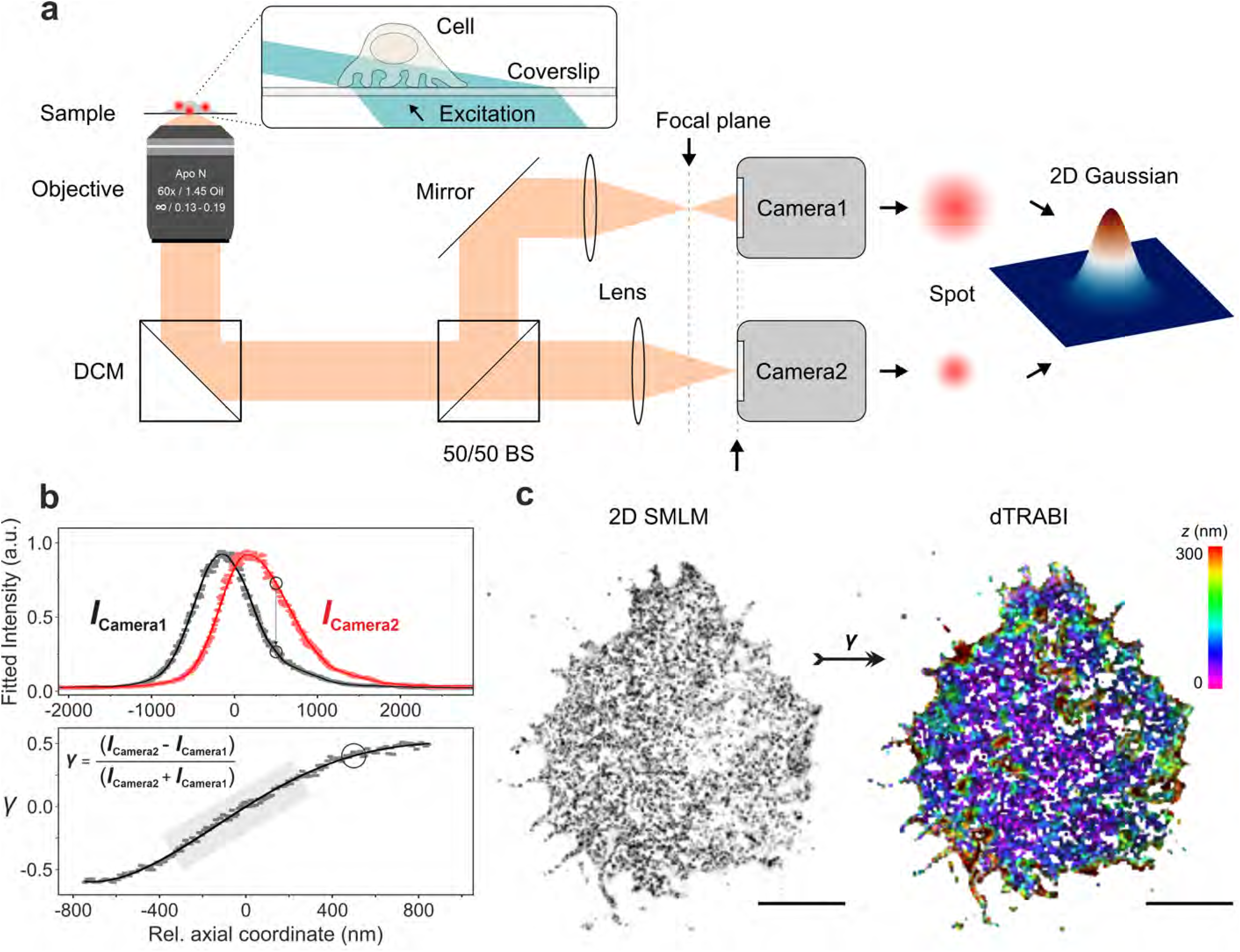
Principle of 3D super-resolution biplane imaging using dTRABI. **a**) Overview of the experimental setup applied to perform 3D dTRABI imaging of T cells. HILO illumination of the sample (blue beam; only shown in enlarged box) triggers fluorescence emission (orange), which is split by a 50/50 non-polarizing beamsplitter (50/50 BS) to acquire biplane images on two separate EM-CCD cameras. The respective imaging lenses are shifted along the optical axis to induce a relative defocus of the image detection on synchronized cameras. Spots, apparent in both detection planes, are fitted by a Gaussian with identically set FWHM. **b)** Using a piezo stage, the focal plane was linearly moved through the sample plane while imaging a single-molecule surface under dSTORM conditions. Hereby, both cameras were synchronized. Fitting the raw PSFs by independent Gaussians with invariable FWHM yielded axially dependent single-molecule intensity curves (upper panel). The relative change of position of the imaging lens in the reflection path is mirrored by the relative shift of the respective intensity curve (indicated by circles). Data points were spline interpolated to guide the eye (solid lines). An axially precise calibration function γ is derived directly from the raw intensities (*I*_*Camera1*_, *I*_*Camera2*_) of corresponding localizations from both cameras as *γ(z)* = *(I* _*Camera2*_ − *I* _*Camera1*_*)(I* _*Camera2*_ + *I* _*Camera1*_)^−*1*^. The running median of the raw data (grey squares) is fitted with a high order polynomial (black line) to generate the basis of the axial lookup table (lower panel). **c**) A two-dimensional high-resolution data set is generated from both image stacks (transmission and reflection path) to create a three-dimensional dTRABI data set according to the calibration. Finally, the transmission localization set is used to render a high-resolution, axially color-coded image of the focused target structure. Scale bars, 5 μm.

#### Intensity-based biplane SMLM imaging - dTRABI

Previously, we reported the temporal, radial-aperture-based intensity estimation (TRABI) method in combination with a biplane detection as a powerful three-dimensional imaging tool with nanometer precision^65,77^. In short, TRABI comprised fitting single-molecule spots by a Gaussian model with an invariant width, a subsequent independent photometric analysis of these spots and a final allocation of the axial coordinate based on a calibration curve. The photometric value was determined from the intensity obtained by the established Gaussian fitting procedure (*I*_Fit_) and background-subtracted reference intensity of the spot (*I*_R_), with *I*_Fit_ < *I*_R_.

Here, we used a simplified but superior version of the TRABI method in which solely the intensity information obtained by the Gaussian fit (*I*_Fit_) was used as metric for both channels in a biplane imaging scheme. Synchronized image stacks of the two channels (transmitted and reflected) generated by dividing the emitted fluorescence equally with a 50/50 beam-splitter, were analyzed by the same Gaussian function with an invariant width for every spot. Spots in the reflected channel were mapped to the corresponding spots in the transmitted channel by the nearest neighbor algorithm (Fig. 2b, see **Methods**). By omitting the standard photometric analysis^65^, the computation time was reduced, and the allowed number of localizations per frame was significantly increased. Therefore, as demonstrated by Fourier Ring Correlation (FRC) of according data sets, we achieve an improved structural resolution while maintaining the same axial localization precision as with the original TRABI approach (Supplementary Figs. S4 and S5, see also **Methods**)^78^. While this resolution-enhancement is more pronounced for structures with a medium or high local signal density, e.g., microtubules, and less distinct, but still apparent, for lower-density structures such as CD4 clusters (Supplementary Fig. S6), there is an additional implication for molecular quantification approaches. We subsequently called the new method dTRABI (for *direct* TRABI; Fig. 2c).

#### Fiducial-free drift and tilt correction

To ensure optimal axial localization over an extended imaging time as well as accounting for subtle sample drift, we developed a fiducial free approach for its correction (see **Methods**). In short, the axial footprint of the entire structure is tracked over time, resulting in a spatio-temporal drift trace (Fig. 3a). This trace is fitted with a high-order polynomial, which serves as the correction term for the raw localizations by linearization. We assumed that in a thin sample layer like the plasma membrane, the spatio-temporal distribution of active photo-switches that reside in their on-state is constant over time. The selection of appropriate regions in axially more extended samples can include structures, where the local z-dimension is restricted. Most data sets that were used to conclude the results in the following paragraphs showed axial drift with different orders of magnitude and non-linearity, which could be accounted for by the described correction approach. However, an additional linear axial tilt was observable in some data sets, most likely due to minor imperfections of the sample holder. Since we assumed the layer closest to the coverslip to be axially flat on the whole-cell scale, we fitted an inclined plane to the raw image, thereby determining its gradient and thus enabling the linearization of the raw localization data (see **Methods** and Fig. 3b).

**Figure 3.**
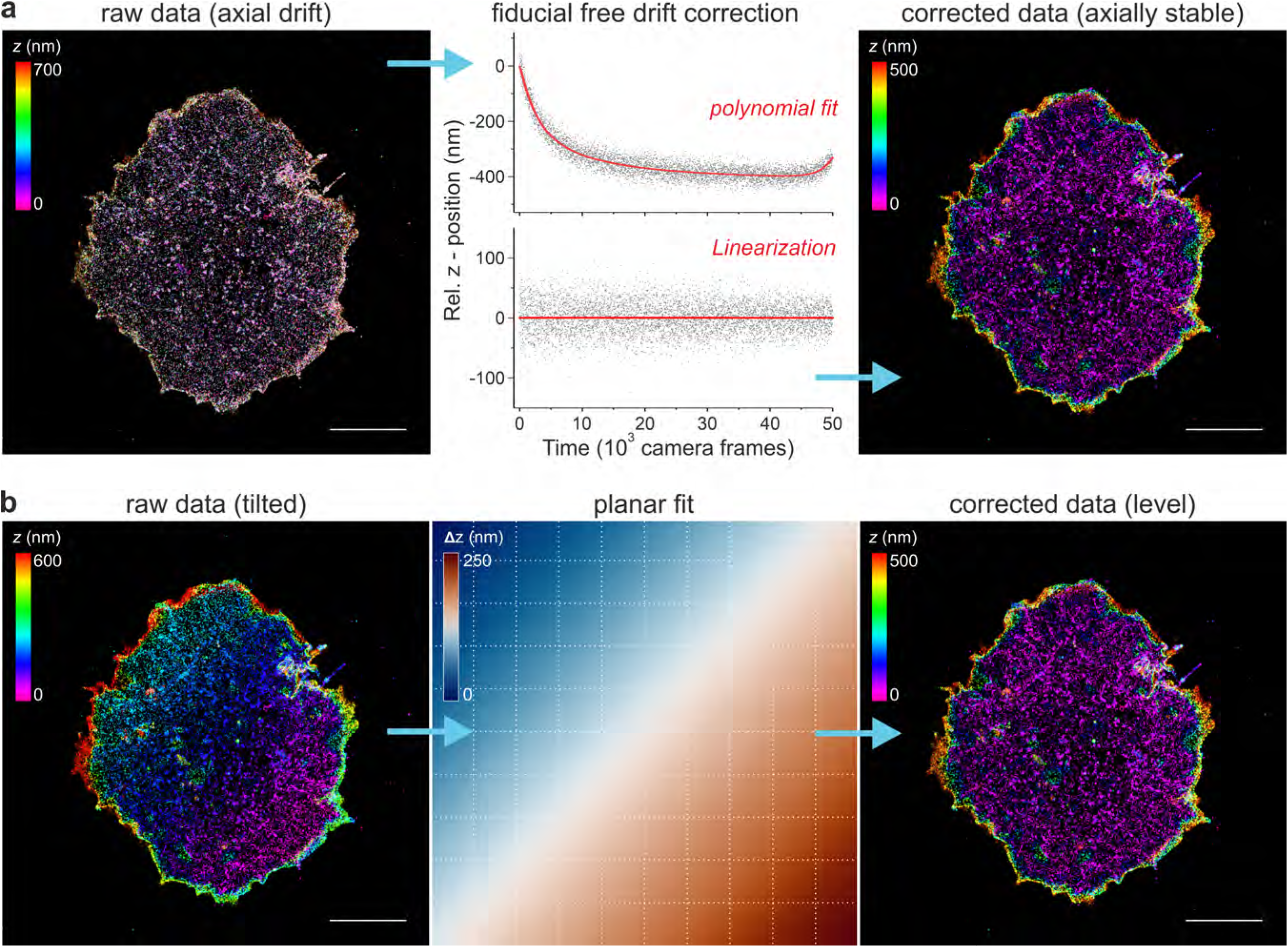
Fiducial free drift and tilt correction of dTRABI data. CD4 WT on Jurkat T cells is shown, where the membrane was directly labelled using Alexa Fluor 647 conjugated OKT-4 antibody. **a)** Principle of the fiducial free correction of axial drift. *Left:* Representative, color-coded high-resolution dTRABI image of an axially unstable sample. Due to the temporal change of the axial coordinate, the resulting image does not exhibit axially distinct features. *Middle:* By extracting the axial localization distribution of the thin membrane layer per frame and tracking it over the entire stack, a spatio-temporal drift trace can be plotted (top). Fitting of the raw data (grey squares) by a high-order polynomial (red line) allows the temporal linearization of the localization data, leading to a stable axial mean value over time (bottom). *Right:* Re-rendering of the drift-corrected localization data reveals a color-coded dTRABI image exhibiting distinct, high-density clusters of CD4 below another, disperse, layer. **b)** Principle of the fiducial free correction of axial tilt. *Left:* Representative, color-coded high-resolution dTRABI image of an axially tilted sample. *Middle:* The axial tilt of the sample is extrapolated by fitting a plane to the raw image. Afterward, the data is linearized by subtracting the local plane-value from the raw localization. *Right:* The tilt-corrected image exhibits a homogeneous color-code in the lowest data layer, indicating no residual axial tilt. Exemplary drift and tilt were simulated based on real data, where the right column represents experimentally corrected real data on which the drift and tilt were projected. Scale bars, 5 μm.

#### Quantitative axial localization analysis

For the analysis of the axial receptor distribution, localization files were loaded and processed in Fiji with custom written scripts (see **Methods**)^79,80^. The analysis comprised three steps: generation of quantitative image stacks, segmentation in regions of interest (ROIs) and axial quantification of individual ROIs. (i) A quantitative 3D image stack (z-stack) was generated with 20 nm pixel size in *x*, *y* and *z*. (ii) These stacks were then segmented by automatically generating multiple regions of interest (ROIs) with an area of 4 μm² within the interior of the cell (Supplementary Fig. S6). (iii) Then, the axial distribution of localizations was analyzed by counting all localizations per ROI along the z-stack. For each ROI, the resulting distribution was fitted to a superposition of two Gaussians (Supplementary Fig. S6). By analyzing the properties of these fits, we acquired parameters that provide quantitative information about the axial distribution of surface receptors (see **Methods**).

### Imaging CD45 nano-topography using the dTRABI approach

To demonstrate the applicability of dTRABI, we labelled CD45 receptors, which are highly expressed on the surface of T cells (> 100.000 copies/cell; ref.^81^) with Alexa Fluor 647-conjugated antibody (MEM-28) and imaged as described above. Cells were immobilized on glycine-coated coverslips to preserve cell surface morphology. Resulting three-dimensional images with color-coded z-axis represent a footprint of a cell on a coverslip (Fig. 4a). The apparent optical depth, which varies between cells (300–650 nm), depends on the available structure in the individual field of view. CD45-labelled cells exhibited a variety of features that can extend far from the coverslip. However, the entire extent of the basal membrane resting on the surface was captured in all displayed and analyzed cells. Fig. 4b shows the three-dimensional image of CD45 which indicates a complex morphology of the cell surface. Magnified images and the corresponding *xz*-projections (Fig. 4c) demonstrate the ability of dTRABI to visualize large membrane protrusions at the edge of the cell (ROIs 1 and 3; blue arrowheads), as well as more subtle membrane extensions towards the exterior (ROIs 1-4; magenta arrowheads) and interior (ROIs 3 and 5; green arrowheads) of the cell.

**Figure 4.**
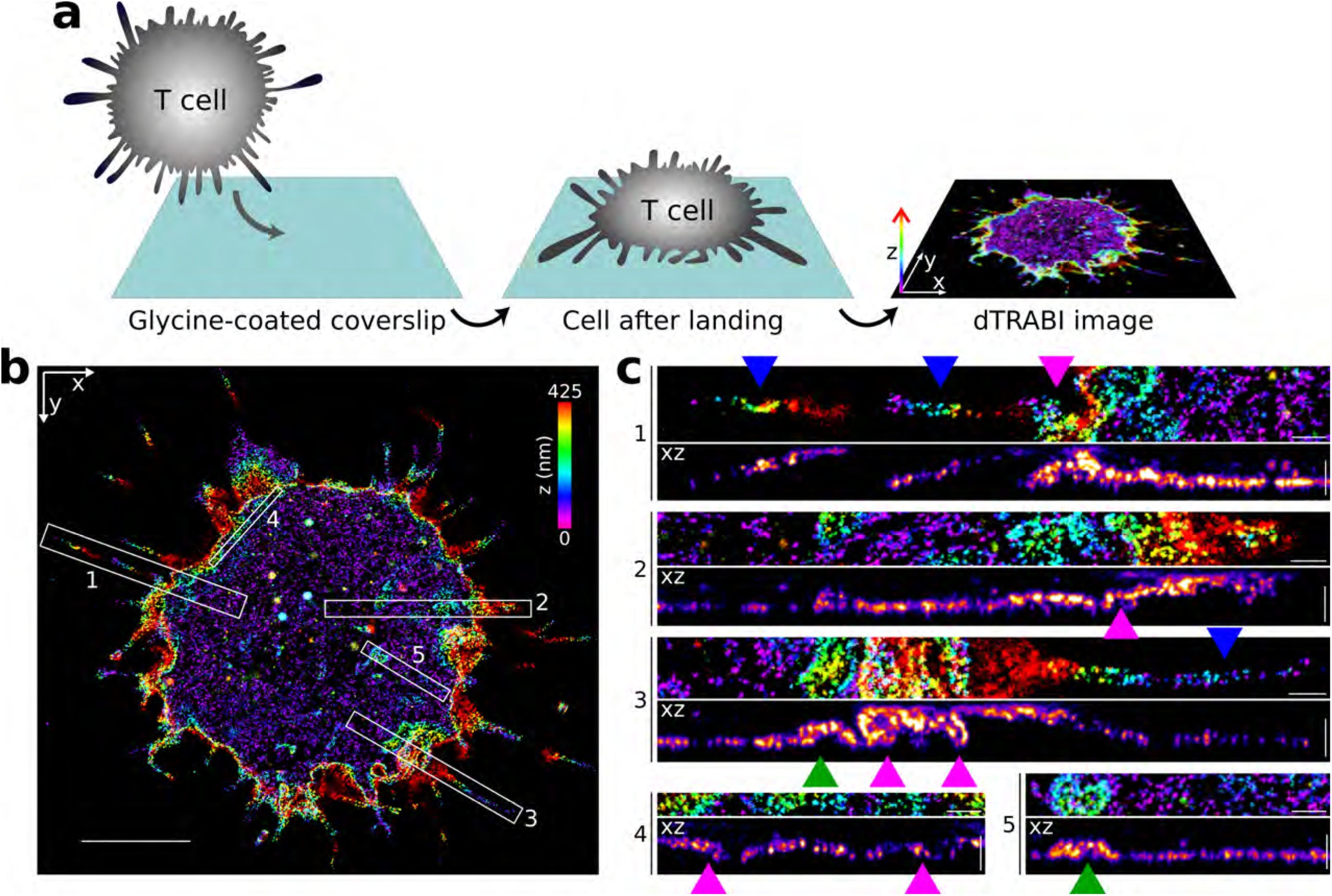
Cell surface receptor nanotopography visualized by 3D TRABI imaging. **a**) Schematic illustration of sample preparation for a receptor nanotopography imaging using dTRABI. A cell in suspension is first immobilized on a glycine-coated coverslip and imaged using dTRABI approach. The resulting three-dimensional dTRABI image represents a footprint of a cell on the optical surface with the color-coded axial position of localizations (right panel). **b)** Three-dimensional dTRABI image of T-cell surface receptor CD45 with selected ROIs exhibiting a broad z-distribution of receptor localizations. **c)** Magnified x-z projections of ROIs 1-5 as in a). Blue arrowheads point to microscopic membrane protrusions at the cell edges, magenta arrowheads to folded nanoscopic protrusions under the cell body and green arrowhead to membrane invagination. Scale bars, 5 μm in b) and 500 nm in c).

### The origin of CD4 microclusters on the surface of unstimulated T cells

Next, we aimed at examining the origin of previously reported receptor (signaling) microclusters^1,4,5^. We recently showed that CD4 receptors accumulate in high-density regions on unstimulated T cells using two-dimensional SR imaging^28^, while the three-dimensional organization of these clusters remained unclear. First, we confirmed that CD4 accumulates in high-density regions on T cells immobilized on glycine-coated coverslips using 2D *d*STORM (Supplementary Fig. S7). Next, high-resolution three-dimensional maps of the CD4 surface distribution were acquired by analyzing the same data sets with the dTRABI approach (Fig. 5a). The high-precision, nanoscopic three-dimensional view of wild-type CD4 indicates that the receptor is clustered and preferentially localized to one topographical level of the membrane (Fig. 5a). This is also demonstrated by plotting the relative z-position of CD4 molecules in the central area, which avoids cell edges (Fig. 5b). On the contrary, a broad surface distribution is evident from the three-dimensional view of a CD4 variant, which cannot be palmitoylated and was previously shown to exhibit random distribution in such unstimulated T cells (CD4 CS1; Fig. 5c-d; ref.^28^). Though these plots provide useful information about axial distribution of receptors in individual cells, their utility for the global distribution analysis is limited by a variation in surface complexity of individual cells. Therefore, the propensity of CD4 molecules to accumulate at specific topographical levels was analyzed in 2×2 μm^2^ square regions of interest (ROIs) selected to cover the cell-coverslip contact area over all tested cells (in total 846 CD4 WT ROIs in 21 cells and 1044 CD4 CS1 ROIs in in 18 cells; Supplementary Fig. S6, Table 1). As mentioned above, for quantitative analyses, the distribution of relative z-positions for localizations in ROIs were fitted with a sum of two Gaussians (bi-Gaussian, Fig. 5e). From these fits, three parameters were derived, i.e., the spread of the bi-Gaussian termed z-distribution width (z_w_) (Fig. 5f-g), distance of its mean values or peak-to-peak distance (p-p) (Supplementary Fig. S8) and the difference of the widths of the two Gaussians (Δ_FWHM_) (Supplementary Fig. S8). By compiling the z_w_ values of all ROIs, a histogram was obtained (Fig. 5f) and further analyzed (**Methods**). For wild-type CD4, we calculated an axial mean value of 120 nm and a standard deviation (s.d.) of 25 nm (cf. Table 1), signifying a strong tendency of wild-type CD4 to accumulate at one specific topographical level in resting T cells (Fig. 5f-g). On the contrary, a lack of palmitoylation in CD4 CS1 variant caused much broader distribution of this receptor on the T-cell surface, with z_w_ predominantly at 141 nm ± 27 nm (mean ± s.d.) and a smaller, yet broader fraction at 247 ± 83 nm (mean ± s.d.). Furthermore, the axial analysis of localizations revealed a more pronounced mean value of Δ_FWHM_ for CD4 CS1 (88 nm) than for CD4 WT (57 nm) (Supplementary Fig. S8c, Table 1).

**Figure 5.**
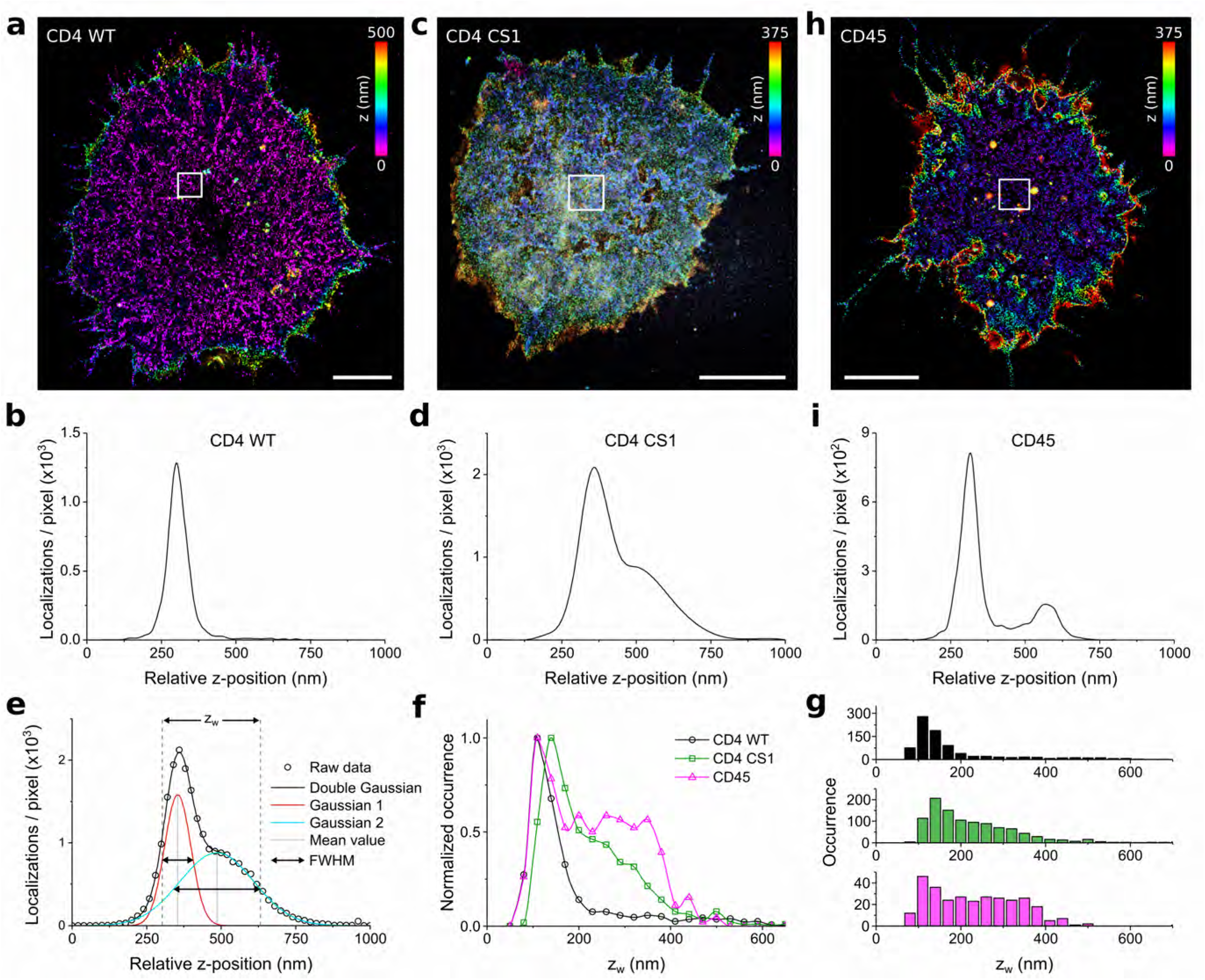
CD4 clusters represent the receptor accumulation at the tips of membrane protrusions. **a)** Representative three-dimensional dTRABI image of CD4 WT at the surface of T cell immobilized on a glycine-coated coverslip. **b)** The axial distribution of CD4 WT localizations of the 4 μm^2^ region of interest (ROI) of the cell in a). **c**) Representative 3D dTRABI image of non-palmitoylatable CD4 CS1 mutant at the surface of T cell immobilized on a glycine-coated coverslip. **d**) The axial distribution of CD4 CS1 localizations of the 4 μm^2^ ROI of the cell in c). **e-g)** Quantitative analysis of the receptor axial (z-axis) distribution on the surface of T cells. Receptors were analyzed by using a bi-Gaussian fit to the axial distribution of localizations for each ROI as in b, d and i (see Supplementary Fig. S6 for more examples) and the FWHM range of the two Gaussian functions represent the z-distribution width (z_w_) as depicted in **e**. Black circles in **e** represent the axial distribution of receptor localizations for a selected ROI, black line the bi-Gaussian fit, which is the sum of two Gaussians as depicted in red and blue, dashed lines in gray depict z_w_ and lines in light grey depict mean values of the Gaussian distributions. The graphs in **f** and **g** represent histograms of z_w_ obtained from 21 CD4 WT cells with 846 ROIs (black), 18 CD4 CS1 cells with 1044 ROIs (green) and 13 CD45 WT cells with 305 ROIs (magenta). The histograms in f) and g) show relative and absolute occurrence, respectively. Data points in b), d), f) and i) were spline interpolated to guide the eye. **h**) Representative 3D dTRABI image of CD45 at the surface of T cell immobilized on a glycine-coated coverslip. Color-bars in the upper right corner a), c) and h) indicate the axial position of the localizations in the image. **i**) The axial distribution of CD45 localizations of the 4 μm^2^ ROI of the cell in h). Scale bars in a) and c) and h), 5 μm.

**Table 1.** Overview of distribution parameters derived from axial receptor analysis. *N* is the number of cells or ROIs. z_w_ = z-distribution width, p-p = peak-to-peak distance, Δ_FWHM_ = width difference; μ is the mean and σ is the standard deviation of the histogram, range = μ ± FWHM/2 with FWHM = 2.355σ; τ is the mean of the exponential distribution. The raw data are shown in Fig. 5 and Supplementary Fig. 8.

|  |  | **WT** | **CS1** | **CD45** |
| --- | --- | --- | --- | --- |
| *N* cells |  | 24 | 18 | 13 |
| *N* ROIs |  | 846 | 1044 | 305 |
| $z_w$ | $\mu_1$ (nm) | 120 | 141 | 118 |
| | $\sigma_1$ (nm) | 25 | 27 | 20 |
|  | range<sub>1</sub> (nm) | 90-150 | 110-173 | 94-145 |
| | $\mu_2$ (nm) | - | 247 | 263 |
| | $\sigma_2$ (nm) | - | 83 | 117 |
|  | range<sub>2</sub> (nm) | - | 149-345 | 126-400 |
| p-p | $\tau$ (nm) | 23 | 36 | 121 |
| $\Delta_{FWHM}$ | $\mu$ (nm) | 57 | 88 | 63 |
| | $\sigma$ (nm) | 15 | 28 | 22 |
|  | range (nm) | 40-75 | 55-120 | 37-89 |

The specificity of CD4 receptor clusters was further emphasized by a less constrained lateral and axial distribution of CD45 in T cells imaged using dTRABI (Fig. 5h,i; 13 cells and 305 ROIs were analyzed). Quantitative analyses were performed consistent with the CD4 variants and demonstrated a broader axial distribution of CD45 receptors than CD4 WT (Fig. 5f-g). We found the z_w_ values to be similarly distributed at 118 ± 20 nm (mean ± s.d.), but in contrast to CD4 WT, CD45 had an additional large fraction of z_w_ at 263 ± 117 nm (mean ± s.d.). This broad CD45 distribution was further noticeable when analyzing the peak-to-peak distance (p-p) of the fitted bimodal distribution to the relative z-position of the receptor (Supplementary Fig. S8b, see **Methods**). The obtained p-p values were exponentially distributed and the mean values for CD4 WT and CS1 were 23 and 36 nm, respectively, whereas the mean value for CD45 was 121 nm. More examples of three-dimensional dTRABI images and their quantitative analysis for all three receptors are shown in Supplementary Fig. S9 and Supplementary Fig. S6, respectively.

### Segregation of CD45 from CD4 clusters in non-stimulated T cells

When analyzing dTRABI images, we noted areas with very few CD45 localizations (Fig. 6a-b; yellow arrowheads). Similarly, CD45 exclusion zones were detectable in two-dimensional SOFI images of antibody-labeled T cells (Fig. 6c). The narrow shape of these exclusion zones was reminiscent of CD4 clusters observed on T cells imaged by the dTRABI approach (Fig. 5a and Supplementary Fig. S9). We thus wondered whether the CD45 exclusion zones represent areas with CD4 accumulation. T cells expressing CD4 WT fused to photo-switchable protein mEos2 (ref.^28^) were labelled with specific anti-CD45 antibodies and analyzed using two-color SMLM. Our data indicate that CD4 and CD45 are essentially segregated into two separate zones on the surface of non-stimulated T cells immobilized on glycine-coated coverslips (Fig. 7a,b). Since CD4 clusters to one topographical level near the T cell-coverslip contact sites (Fig. 5a,b,f), we suspect that CD4 preferentially accumulates at the tips of membrane protrusions (e.g., microvilli) and CD45 to the shaft of these structures and their base at the plasma membrane (Fig. 7c). Indeed, we observed the accumulation of CD4 at the tips of large T-cell membrane protrusions in several cells (Fig. 7d,e). In turn, the shaft of protrusions was extensively covered with CD45 signal. Often, CD45 was essentially segregated from CD4 in these nanoscopic membrane structures (Fig. 7e). These data are in agreement with recently reported observations that CD45 segregates from microvilli tips in resting and activated T cells^82^.

**Figure 6.**
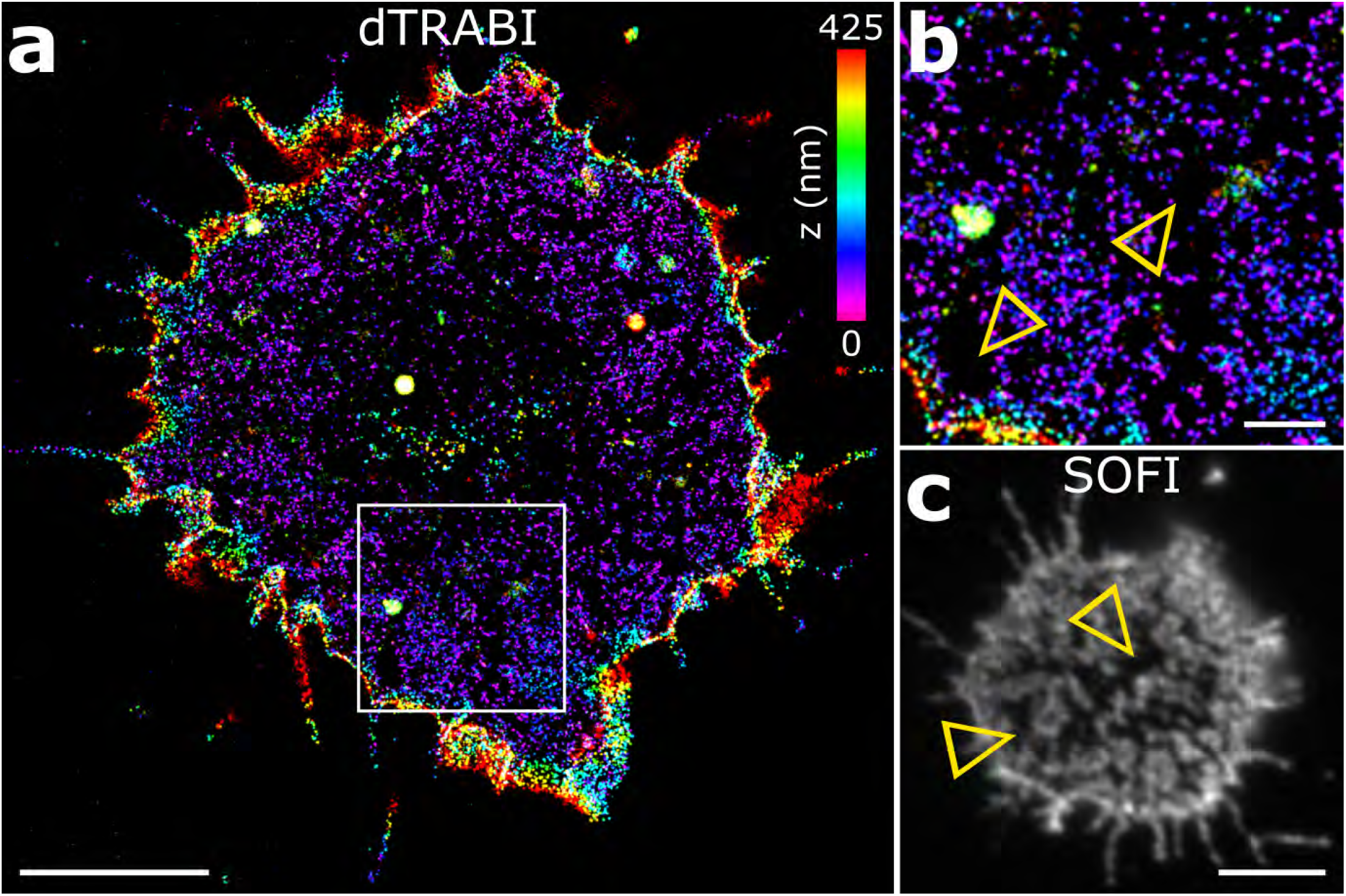
Areas on the surface of resting T cells lacking CD45. **a**) Three-dimensional dTRABI image of CD45 on T cell. **b**) Magnified ROI as in a) with indicated areas lacking CD45 localizations (yellow arrowheads). **c)** 2D SOFI image of CD45 on T cell immobilized on a glycine-coated coverslip. CD45 was labelled using Alexa Fluor-647 conjugated MEM-28 antibody. Yellow arrowheads indicate cell surface areas lacking CD45 signal. Scale bars, a) 5 μm, b) 1 μm and c) 5 μm.

**Figure 7.**
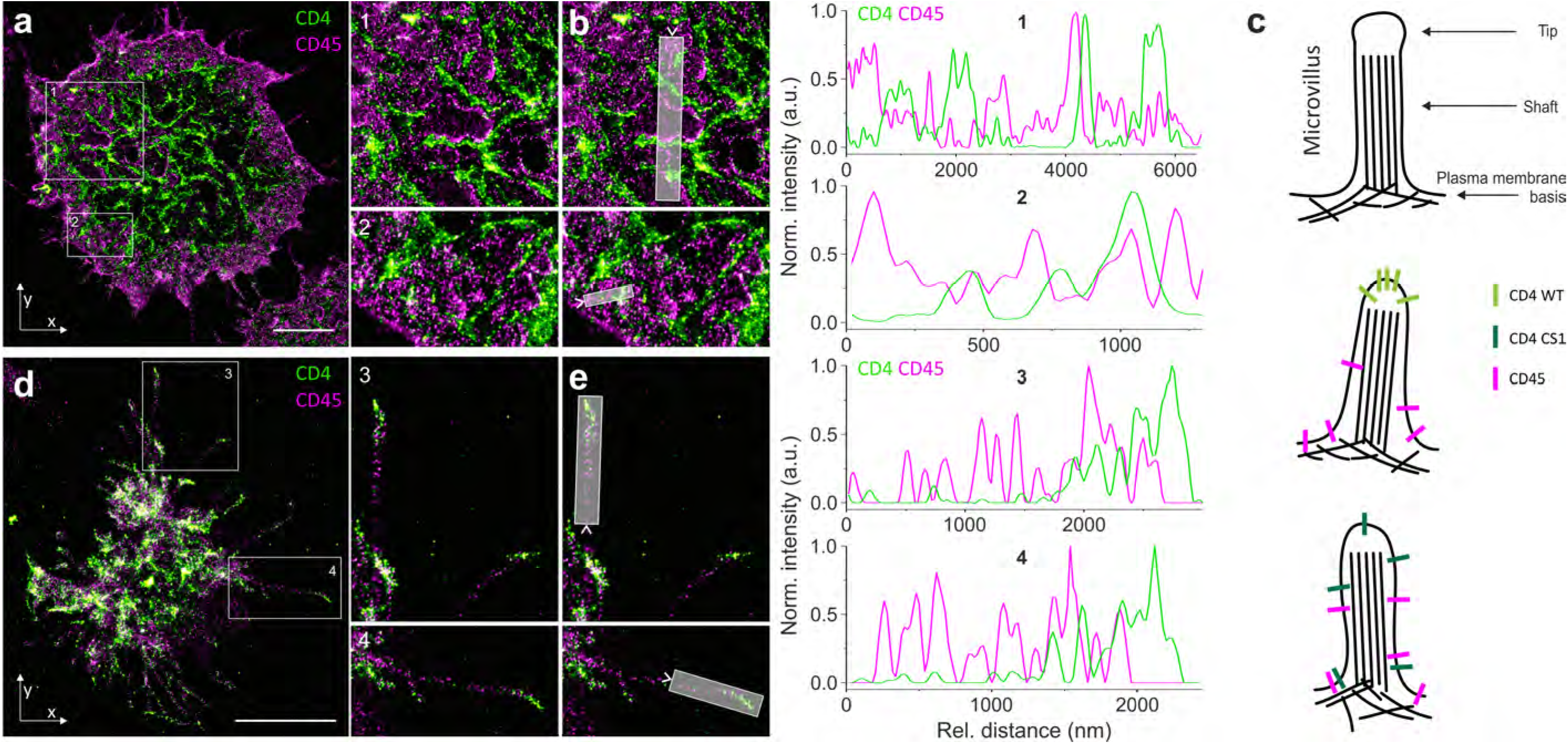
Nanoscopic segregation of CD4 WT and CD45 on the surface of resting T cells. **a**) Representative two-dimensional image of T cell surface sequentially analyzed for CD4 WT (green) and CD45 (magenta) by SMLM. CD4 was visualized as mEos2 fusion protein (PALM) after transient transfection of cells, and the surface CD45 was labelled using Alexa Fluor-647 conjugated MEM-28 antibody (dSTORM). ROIs 1-2 were zoomed to indicate details of proteins’ distribution (right side). **b**) Intensity line-profiles were measured along the transparent grey regions indicated in ROIs 1-2 (as in a). Green line represents CD4 WT and magenta CD45 signals. **c**) Schematic illustration of microvillus with indicated structural segments: tip, shaft and the basis. The two panels below indicate the organization of CD4 and CD45 on the protrusions in cells expressing native CD4 (WT; middle panel) or its non-palmitoylatable variant (CS1; lower panel) CD4 as indicated by the nanoscopy. **d)** Two-dimensional SMLM image of a selected T cell captured during its association with the optical surface which was sequentially analyzed for CD4 WT and CD45 (as in a). The accumulation of CD4 WT on the tips of large membrane protrusions was observed in 10 out of 32 imaged T cells. ROIs 3-4 were zoomed to show the details of membrane protrusions with accumulated CD4 WT on their tips (right side). **e**) Intensity line-profiles were measured along the transparent grey regions indicated in ROIs 3-4. The arrows indicate the onset of line-profiles. Scale bars, 5 μm. Images from three independent experiments are shown (n=32).

## DISCUSSION

In this work, we have introduced a new approach for molecular mapping of the membrane receptor topography with nanometer precision. The method combines the use of glycine-coated optical surfaces, which preserve the native membrane morphology of immobilized cells with an advanced and simplified dTRABI algorithm for quantitative, near isotropic three-dimensional SMLM imaging. Using directly labelled antibodies and dTRABI processing of biplane SMLM data, we achieved 10–20 nm localization precision in all spatial directions. This enabled us to reveal the origin of CD4 clusters and the involvement of cell surface morphology in CD45 segregation from signaling receptor clusters.

Our principal task was to preserve fine structures on the surface of T cells and to analyze the distribution of signaling receptors therein. Even though flat in most common illustrations, the plasma membrane of differentiated metazoan cells is a three-dimensional structure densely covered with finger-like protrusions (microvilli, filopodia), ruffles and invaginations (endocytic cavities and podosomes)^40^. Membrane imaging with two-dimensional methods thus leads to simplifications and, potentially, misinterpretations. The availability of SR microscopy capable of resolving nanoscopic three-dimensional structures is currently limited to specialized nanophotonics laboratories. Nevertheless, standard methods of sample preparation for imaging (e.g., on PLL-coated coverslips) rapidly damage the fragile surface morphology of studied cells, as indicated also by our data showing that the T-cell surface complexity is diminished within a few minutes of the first contact with a PLL-coated coverslip. Several materials were previously developed to prevent cell deformation during their immobilization in a three-dimensional space, for example, Matrigel and synthetic hydrogels. Matrigel enables three-dimensional cell cultures, but it is derived from the extracellular matrix of mouse tumors and exhibits autofluorescence. On the contrary, fluorescence properties of synthetic hydrogels (e.g., CyGEL Sustain) can be controlled and the cells prepared in hydrogels display numerous protrusions on their surface, including nanoscopic structures that require SR microscopy for their detection^58^. However, TIRF and HILO sample illumination techniques which can reduce the out-of-focus background signal and, thus, improve the quality of SMLM images, enable fluorescence signal detection solely in the vicinity of the optical surface^76,83^. Hydrogels do not allow efficient positioning of immobilized cells at the optical glass.

Here, we developed and analyzed glycine coating of coverslips, which better preserves cell membrane morphology. The coating of coverslips with a narrow layer of gel-like glycine improves the stability of membrane protrusions and facilitates the positioning of immobilized cells close to the optical surface. Importantly, cells immobilized on glycine exhibit improved viability and reduced non-specific stimulation compared to PLL-immobilized cells. Of note, the preparation of glycine-coated coverslips is straightforward, fast (<30 min) and inexpensive. All these properties qualify glycine-coating of coverslips for the advanced imaging of cells, including high-resolution three-dimensional dTRABI mapping of cell surface molecules.

We have also developed the dTRABI algorithm to allow for faster and computationally more efficient processing of SMLM data, while achieving significantly higher structural resolution, than that found with our original TRABI approach. It is built for a quantitative analysis of the axial distribution of fluorescent molecules. Importantly, dTRABI can be used with any localization software that supports fixed Gaussian fitting in combination with a biplane detection scheme. In contrast to classical biplane imaging that compares the width of spots evaluated by free Gaussian PSF fitting, dTRABI employs fixed-width Gaussian fit which facilitates an improved localization precision. The ability of dTRABI to localize fluorescent molecules on complex, irregular, three-dimensional cellular structures was highlighted on a T cell that was not included in the quantitative analysis of an axial receptor distribution (Supplementary Fig. S10). Complex membrane structures formed by this dying cell were effectively labeled and visualized using the dTRABI approach.

Importantly, dTRABI offers a tool for mapping localization of molecules within ~1 μm of the optical surface. Such working space provides a sufficient axial signal penetration to visualize the eukaryotic plasma membrane with its nanotopography. Under the conditions used in our experiments, the plasma membrane protrusions were positioned 100–400 nm from the optical surface (Fig. 5). Thus, molecules that do not localize to the protrusions can be monitored as well. This was facilitated by the folding of membrane protrusions, the length of which often exceeded the axial depth of our method, under the cell body (Fig. 4 and Supplementary Fig. 11a,b). Evidently, a partial loss of molecular localizations in the areas where the base of the plasma membrane is positioned outside of the optical limit of this method is caused by a diverse positioning of cells and intensive bending of cell-glass contacts (Supplementary Figs. S9 and S11c,d).

The quantitative character of dTRABI data also enabled a detailed analysis of axial receptor distribution, which we modelled by fitting a sum of two Gaussian functions (bi-Gaussian) to the axial localization data. Although CD4 distribution can be modelled by a single Gaussian to a high degree of confidence, a bi-Gaussian is even more accurate and accounts for a slight asymmetric organization of receptors. In contrast, non-palmitoylatable CD4 CS1 mutant does not follow a mono-Gaussian distribution and the bi-Gaussian is essential. The analysis allowed us to derive quantitative parameters describing axial receptor distribution in detail, i.e., z_w_, p-p and Δ_FWHM_. All these analytical approaches emphasized the narrow and symmetrical distribution of CD4 WT, while CD4 CS1 and CD45 show significantly broader distributions.

To highlight the effectiveness of our new imaging approach, we explored two important biological questions: i) what is the origin of receptor microclusters in T cells, and ii) how does membrane morphology affect segregation of signaling receptors into different areas of the plasma membrane? Both questions cannot be studied by previously reported protocols. Using quantitative SOFI analysis, we have previously shown that CD4 accumulates in high-density regions (clusters) in unstimulated T cells by a process that depends on its post-translational lipid modification, palmitoylation^28^. However, as in many other cases, these data represented two-dimensional projections of receptor localizations on the complex T-cell surface. Using dTRABI imaging of unstimulated T cells immobilized on glycine-coated coverslips, we demonstrate that CD4 accumulates on the tips of microvilli. Thus, CD4 clusters, and potentially other clusters, represent receptors (or other molecules) trapped at the tips of membrane protrusions, stressing that intact cell surface morphology is essential for the proper spatial representation of signaling molecules. Our data are in agreement with recently reported clustering of CD4 on the tips of T cell microvilli ^84^ but provide more detailed and quantitative insight into the CD4 distribution to these structures. Importantly, the mutant CD4 CS1 variant which cannot be palmitoylated distributed more randomly over the complex surface of T cells. Targeting to or stabilizing proteins on membrane protrusions thus may be another, previously unreported role of protein palmitoylation^85^.

The best-described function of CD4 is to deliver critical Lck kinase to the proximity of T cell receptor for a rapid response to antigens^60^. TCR was recently reported to accumulate in T cell microvilli by projecting a two-dimensional SMLM image onto the morphological map generated by varying-angle TIRF microscopy (VA-TIRFM) of a membrane dye ^84,86^. VA-TIRFM is diffraction-limited and, therefore, does not provide a high-resolution map of cell surface nanotopography. However, the data indicated the accumulation of TCR in domains formed by or dependent on membrane morphology. We thus speculate that CD4 and TCR localize to the protrusions, the tips of which can represent membrane areas with concentrated signaling molecules arranged to rapidly respond to appropriate signals (Fig. 7c). How these molecules accumulate in these structures and whether the tips of protrusions undergo further compartmentalization remains unclear. In future ventures, we will establish a robust workflow consisting of reliable sample preparation, labelling and multi-color dTRABI analysis to investigate the multi-molecular architecture of these structures. Our current approach already provides a good basis for such advancement and is adaptable to multiple avenues of inquiry.

Several molecules, including CD45, were shown to segregate from signaling microclusters (e.g., refs.^63,87^). Using dTRABI and glycine-coated coverslips, we demonstrated that CD45 distributes over a broader axial spectrum than CD4. Since CD45 segregates from CD4 in non-stimulated T cells under the experimental conditions used in this study, we conclude that CD45 segregates from signaling molecules by localizing to the shaft of microvilli and basis of the plasma membrane (Fig. 7c). The molecular mechanism of such segregation remains unknown. Even though listed in a high-throughput study of palmitoylated proteins^88^, direct experimental proof of CD45 palmitoylation is missing. Since the non-palmitoylated CD4 CS1 variant exhibits a more random distribution compared to its native variant, we speculate that the lack of CD45 palmitoylation contributes to its preference for the basis of the plasma membrane.

In summary, we provide a new and undemanding workflow to study nanotopography of receptors at the cell surface. We highlight the importance of appropriate sample preparation for the imaging of three-dimensional structures. The improved algorithm of dTRABI exemplifies an easily implemented and high-quality method to generate three-dimensional SMLM data. Finally, we demonstrate the applicability of the workflow by answering two critical questions related to the localization of surface receptors on lymphocytes. Even though the method was used here for human receptors, it can be implemented for molecular characterization of the surface on other organisms (e.g., yeast, plants) but also for complex nanomaterials that do not exceed the current axial penetration of dTRABI (~ 1 μm).

## METHODS

### Cell culture and transfection

Jurkat T cells (clone E6; ATCC) and their CD4-knock-out variants (Jurkat CD4-KO cells) were grown in RPMI-1640 medium (Gibco) supplemented with 10% fetal bovine serum (Gibco), Non-essential amino-acids (Gibco) and 10mM HEPES under controlled conditions in a humidified incubator at 37°C, and 5 % CO_2_ (Eppendorf). Jurkat CD4-KO cells were derived from wild-type Jurkat T cells (clone E6; ATCC) using CRISPR/Cas9 technology as described^89^. Both cell lines were regularly tested for morphology and mycoplasma infection.

For the expression of exogenous proteins, Jurkat T cells and their CD4 KO variant were transiently transfected with plasmid DNA using Neon^®^ transfection system (Thermo Fisher Scientific) according to the manufacturer’s instructions. Briefly, 1 μg of vector DNA was used per 200 000 cells in 0.5 ml culture medium. The instrument settings were: 3 pulses, each 1350 V for 10 ms. Transfected cells were used within 36 h since the transfection.

### Sample preparation for SR microscopy

#### Cleaning and handling of coverslips and solutions for microscopy

High precision microscopy coverslips (round, 25 mm in diameter; Marienfeld) in Teflon holders (Wash-N-Dry Coverslip Rack; Diversified Biotech Inc.) were cleaned by incubation at 56°C overnight in 2% Hellmanex III (Hellma Analytics) dissolved in ultrapure Milli-Q^®^ water (Millipore) followed by 30 minutes sonication in a heated sonication bath. Several washes in ultrapure water and one additional sonication step were used to remove all traces of Hellmanex III components. Cleaned coverslips were stored in ultrapure water to avoid drying and contamination with particles of a dust from air.

All other glassware was regularly treated with piranha solution (3 parts of 30% hydrogen peroxide and 7 parts of concentrated sulfuric acid) for 45 minutes followed by several washes with ultrapure water. All solutions were made from concentrated stocks stored in glass containers, diluted with ultrapure water and filtered using syringe filters with 0.22 μm pores (TPP) into piranha-treated glassware.

#### Glycine coating of coverslips

Ultraclean coverslips were coated by applying 0.5 ml of 2M glycine solution and incubation for 20 min at room temperature in the laminar flow box. Afterwards, the liquid phase was aspirated, and the hydrogel was washed with ultrapure water. Coverslips coated with glycine hydrogel were used immediately for immobilization of cells. The hydrogel-coated coverslips can be stored at 4°C covered with a layer of ultrapure water for several days.

#### PLL coating of coverslips

Ultraclean coverslips were coated by applying 0.5 ml of 0.01% (w/v) PLL solution in ultrapure water and incubation for 20 min at room temperature in the laminar flow box. Unbound PLL was removed by aspiration of the liquid and a single wash with ultrapure water. Dried, PLL-coated coverslips can be stored for several weeks. For SR imaging, freshly prepared PLL-coated coverslips were used exclusively.

#### Immobilization and fixation of cells for SR imaging

Jurkat T cells express varying levels of CD4 which are changing when cells are grown in culture. Therefore, we used transiently transfected Jurkat-CD4KO cells with reintroduced CD4-GFP to study CD4 surface distribution with dTRABI. Cells expressing similar levels of CD4-GFP were selected for imaging and data processing.

For SR imaging, after centrifugation for 3 minutes at 500 g (room temperature), cells were resuspended in pre-warmed PBS (made from 10x stock, Gibco) and seeded on glycine-coated coverslips immediately after the aspiration of a coating liquid. Cells were enabled to land on the coated optical surface for 10 minutes at 37⁰C in the CO_2_ incubator. Afterwards, cells were fixed with pre-warmed 4% paraformaldehyde (Electron Microscopy Sciences) containing 2% saccharose in PBS for 45 minutes at room temperature and the process was stopped with 50 mM NH_4_Cl (Sigma-Aldrich) in PBS and three rounds of washing with PBS.

#### Immunofluorescence

For labeling, cells were first incubated with 5% BSA in PBS (Blocking solution) for 1 hour at the ambient temperature to prevent non-specific binding of antibodies. Immunostaining of specific receptors was performed by incubating cells overnight with AlexaFluor 647 conjugated primary antibodies (human CD4: OKT4, dilution: 1:100, source: Biolegend; human CD45: MEM-28, dilution: 1:2000, source: ExBio) diluted in Blocking solution. The process was performed at the ambient temperature in a humid chamber to avoid drying of the solutions. After removing of redundant antibodies and 3 times washing with PBS for 5 minutes, the cells were post-fixed with 4% PFA for 5 minutes and washed 5 times with PBS.

The identical protocol was applied for the staining of CD45 on Jurkat CD4 KO cells transfected with pXJ41-CD4-mEOS2 plasmid for 2-color 2D SMLM analysis.

### Single-Molecule Localization Microscopy

Three-dimensional SMLM experiments for dTRABI analysis were performed on a home-built wide-field setup, which is described elsewhere in detail^65^. Raw image stacks were analyzed with rapidSTORM 3.2 (ref.^90^). Herein, the FWHM was set to 300 nm as an invariant parameter. Furthermore, the lower intensity threshold was set to 500 photons and the fit window radius to 1200 nm. All other fit parameters were kept from the default settings in rapidSTORM 3.2. Linear lateral drift correction was applied manually by spatio-temporally aligning distinct structures to themselves. This was facilitated by color-coding of the temporal coordinate with the built-in tool.

Two-color SMLM experiments were performed on a Nikon Eclipse Ti microscope, which is specified elsewhere in detail^44^. Contrary to the case of dTRABI analysis, the FWHM was set as a free fit parameter, but in the limits of 275–750 nm, both for Alexa Fluor 647 antibody and mEOS2 fusion protein localizations. All other parameters were kept consistent to the previous experiments. Prior to imaging, a glass surface with Tetraspeck beads (Thermo Scientific) was imaged with alternating 561 nm and 647 nm excitation to create a nanometer precise map to allow the correction of chromatic shift. The 561 nm excitation channel was then mapped onto the 647 nm after the localization step.

Prior to acquisition, cells were irradiated in epifluorescence illumination mode to turn emitters, which were out-of-focus in the HILO illumination scheme, into the dark state. In all experiments, the length of the acquisition was set to capture the majority of emitters, i.e. imaging was concluded when only a very minor number of active emitters was detectable. Typical acquisition lengths were 60,000–120,000 frames for Alexa Fluor 647 channel and 30,000–60,000 frames for mEos2 channel, where integration times were set to 20 ms and 16 ms in the single- and dual-color cases, respectively. Hereby, mEOS2 was excited at 561 nm and activated with 405 nm. The activation power density was increased over the time to create an almost constant signal density.

### Intensity based biplane imaging – dTRABI

#### The principle of dTRABI

We previously reported TRABI-based biplane (BP-TRABI; ref.^65^), which utilizes the photometric ratios *P* of the molecules for the image reconstruction. *P* was calculated as the quotient of the fit intensity (*I*_*Fit*_) and reference intensity (*I*_*R*_), where *I*_*Fit*_ was derived from a PSF fit with fixed width, which was set in rapi*d*STORM software. For BP-TRABI, the photometric ratio of both planes (*P*_*1,2*_) was calculated according to

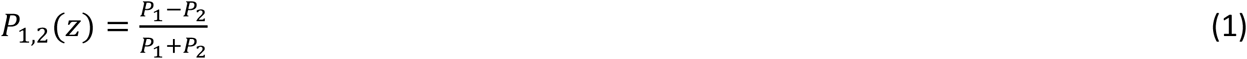

with *P*_*1*_ and *P*_*2*_ being the photometric ratio of the individual planes.

The quality of the resulting axial coordinate depended on the precision of the TRABI intensity measurement. In order to exclude any interference from neighboring emitters as well as temporal overlap with other fluorophores, a set of rigorous exclusion criteria for spots was employed. As a consequence, many localizations given by the localization software were rejected by TRABI. To compensate, significantly longer image stacks had to be recorded to ensure structural consistency in the reconstructed images regarding the Nyquist-criterion. Though this was already an inconvenience for common SMLM organic fluorophores, which exhibit high repetition counts, it can be a significant obstacle for imaging approaches utilizing photo-activatable or -convertible fluorescent proteins. Furthermore, in cases of sub-optimal photo-switching rates that lead to high spot densities, the TRABI approach was computationally expensive or could even fail to produce an image due to a high rejection rate of fluorescent spots.

For a reasonable large TRABI radii *I*_*R*_ converges for both spots in both planes, thus Eq. (1) can be effectively simplified to

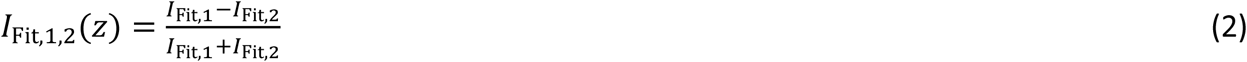

As Eq. (2) demonstrates, in order to achieve three-dimensional BP imaging, there is no need for an actual photometric TRABI analysis anymore, since *I*_*R*_ is cancelled out. Therefore, dTRABI allowed for higher emitter densities, was quicker and required less input parameters.

#### Calibration and allocation

Raw calibration curves were generated by linearly moving the focal plane through the sample plane while imaging a single-molecule surface under dSTORM conditions as previously described^65^. For this, a surface of BSA was doped with BSA molecules, attached to a short DNA sequence, labeled with Cy5. Fitting the raw emission patterns by independent Gaussians with the same fixed FWHM yielded axially dependent single-molecule intensity curves. An axially precise raw calibration function γ was derived according to Eq. (2). The running median (binning width 25 nm) of the raw data was fitted with a high order polynomial to generate the axial lookup table.

After an initial rough alignment between the channels, experimental localizations from both optical channels were assigned by a framewise, linear nearest neighbor analysis. Here, the distance threshold was set to 500 nm, which seemed a reasonable value for the robust allocation between channels in a semi-sparse single-molecule environment. From these sets of localizations, the axially dependent intensity quotient *I*_Fit,1,2_ (Fig. 2b) was calculated and roughly allocated to the look-up table (LUT). The final axial coordinate was determined by a linear interpolation of *I*_Fit,1,2_ between its “left” and “right” nearest neighbor coordinate of the LUT. Obtained axial coordinates were corrected for the refractive index mismatch as previously described^65^.

#### Drift and tilt correction

Since the plasma membrane can be seen as flat over the whole-cell scale, we reckoned that it can be used as its own fiducial marker. We traced the spatio-temporal axial footprint of the entire membrane by fitting the raw localizations by a high-order polynomial in time. This is followed by the straightforward temporal linearization of the localization data, leading to a stable axial mean value over time. In order to instantly assess the quality of three-dimensional SMLM data we suggest looking for white regions in the color-coded image. In our experience, the abundance of these features usually suggests significant axial drift. However, even a subtle axial drift, which cannot be easily recognized by eye, will be detected by the approach described above. Additionally, non-linear drift, commonly occurring due to heating and resting of threads, can be accounted for.

Since we regularly observed a subtle axial tilt in the nanoscopic color-coded images due to the usage of a round coverslip in a magnetic holder, we developed a simple correction workflow (Fig. 3). The axial tilt of the sample is extrapolated by fitting a plane to the raw image data (pixel size 100 nm). Afterwards, the data was linearized by simply subtracting the precise local plane-value from the raw localization.

#### Localization precision calculation

Drift and tilt corrected localizations were tracked in time by determining the three-dimensional nearest neighbor distances of localizations in consecutive frames. Localizations constructing a track with a total inter-localization distance of less than 75 nm were considered to stem from the same fluorophore. For each sample type, we combined the nearest-neighbor tracks from all recorded independent fields of view, calculated the deviation from the mean coordinate of each track for all relevant spatial coordinates and derived a normalized histogram (Supplementary Fig. S6). By fitting these distributions with a Gaussian, we derived the localization precisions as the standard deviation of the mean.

#### Fourier Ring Correlation (FRC) analysis

Standard TRABI-biplane files were created with a TRABI radius of 8.5 camera pixel, a base-jump of 2 frames, an exclusion zone factor of 2, a highlander-filter of 250 frames and a number of averaged background frames of 5. Three-dimensional data sets were then sorted into groups of localizations apparent in even and odd numbered frames, respectively, which were rendered to high-resolved images in rapi*d*STORM with 10 nm pixel size. FRC was performed on these two respective images, using the NanoJ-Squirrel Fiji plugin^78^ with the input of 20 segments per dimension. Since the resulting FRC resolution maps showed a prevalence for extreme outliers, we chose the median FRC resolution over the entire displayed image as resolution criteria (Supplementary Fig. S5).

### Axial localization distribution analysis

Localization files obtained from rapi*d*STORM were loaded and processed in Fiji ^79^ with custom written scripts^80^. First, all localization coordinates in *x, y* and *z* were used to generate a quantitative stack of 2D images with 20 nm pixel size using a separation of 20 nm in *z* (i.e. z-stack). The numeric value in each pixel was equivalent to the number of localizations. Then the z-stack was segmented into defined regions of interest (ROIs). The interior of the cells was manually selected to serve as boundary ROI, in which the edges of the cells were spared. Within this master ROI a set of squared ROIs with 2×2 μm^2^ area was automatically generated. The ROIs were allowed to be confined by the border of the master ROI and only kept when more than 75% of the ROI area, i.e. > 3 μm were preserved (Supplementary Fig. S6).

Afterwards, the localization density of each ROI was analyzed by accumulating the grey values per ROI within the image stack, i.e. stepwise in *z* every 20 nm, and plotted as a function of *z*. The plot was then fitted to a bi-Gaussian function of the form

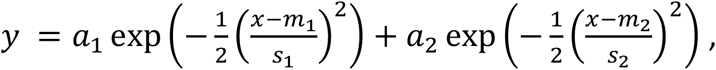

with *a*_1,2_ as amplitude, *m*_1,2_ as mean value and *s*_1,2_ as standard deviation of each Gaussian.

To characterize the axial distribution of localizations of CD4-WT, CD4-CS1 and CD45, three parameters were derived from this fit. 1) the z-distribution width (Fig. 5e-g) calculated according to

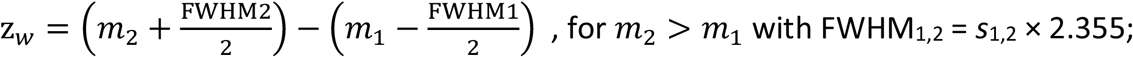

2) the peak-to-peak distance (Supplementary Fig. S10) according to p-p = |m_2_ − m_1_|; and 3) the width difference of the two Gaussians (Supplementary Fig. S8) according to Δ_FWHM_= |FWHM_1_ − FWHM_2_|.

### Quantifying axial receptor distribution

For quantifying the obtained axial receptor distribution in Fig. 5f, we fitted the histogram of z_w_ for CD4 WT, CD4 CS1 and CD45 with a model of Gaussian functions. CD4 WT was fitted using a mono-Gaussian and yielded the mean value μ and standard deviation σ, the range of z_w_ in Table 1 was stated as μ ± FWHM/2, with FWHM as full width at half maximum of the distribution. CD4 CS1 and CD45 z_w_ histograms were fitted using a bi-Gaussian yielding μ_1_, μ_2_ and σ_1_, σ _2_ as mean and standard deviation, respectively. The range of values in Table 1 was stated as μ ± FWHM/2. The p-p distributions (Supplementary Fig. S8) were modeled with a mono-exponential decay, and the inverse of the decay constant of the fit (τ), i.e. the value at which the amplitude is reduced to 36.8%, was stated as mean value. The distribution of Δ_FWHM_ was modeled with a mono-Gaussian, thus obtaining μ and σ as mean and standard deviation, respectively; the range of values in Table 1 was stated as μ ± FWHM/2.

### SOFI

Sample preparation, coverslip coating and image acquisition were performed as described above for SMLM imaging. The raw data were analyzed using balanced SOFI algorithm as described before^28,91^.

### Calcium measurements

#### Calcium sensor

Plasmid DNA with the ultrafast, genetically encoded calcium sensor GCaMP6f_u_ was a kind gift from Katalin Török and Silke Kerruth (St George’s, University of London). The coding sequence was amplified using primers AATAGATCTGCCACCATGGGCTGCGTGTGCTCCTC and AATGGATCCTCACTTCGCTGTCATCATTTGTACAAA and subcloned into pXJ41 vector using *BglII* and *BamHI* restriction sites as described before^92^.

#### Sample preparation

For calcium measurements, the coverslips were coated with 0.01% (w/v) poly-*L*-lysine (Sigma-Aldrich) or 2M glycine as described above. For coating with OKT3 (anti-CD3ε) antibody, clean coverslips were incubated with 0.01 μg/μl OKT3 in PBS for 30 minutes at 37°C. After a brief wash with PBS, the coverslips were used within a day. Prior to acquisition, coated coverslips were mounted into a ChamLide holder (Live Cell Instruments), filled with 500ul of color-free medium, placed on the microscope and focused to the focal plane. The measurements were performed twenty hours after transfection of Jukrat T cells with the calcium sensor. For image analysis, cells were washed with PBS, resuspended in phenol red-free RPMI-1640 media (Sigma-Aldrich) supplemented with 2mM *L*-glutamine, 10mM HEPES, 1mM CaCl2 and 1mM MgCl_2_ and dropped onto the prepared coverslip while running the image acquisition.

#### The microscope setup and image acquisition

Live cell calcium mobilization imaging was performed on a home built TIRF microscope consisting of the IX73 frame (Olympus), UApo N, 100x 1.49 Oil immersion TIRF objective (Olympus) and OptoSplit II image splitter (CAIRN Optics) mounted on the camera port. Samples were illuminated using 200mW 488 nm laser (Sapphire, Coherent) in a TIRF mode and the intensity was regulated by acousto-optic tunable filter (AOTFnC-400.650-TN, AA Optoelectronics). Fluorescence emission was detected by an EMCCD camera (iXon ULTRA DU-897U, Andor) with EM gain set to 200. Images were taken in 500ms intervals with the exposure time set to 50 ms.

#### Data processing

Calcium mobilization was quantified by calculating mean fluorescence in cells immobilized on coated coverslips over the period of 10 minutes. Fiji/ImageJ software was used for the analysis^79^. Graphs in Supplementary Fig. S3 represent a mean fluorescence changes in cells immobilized on PLL-, glycine- or OKT3-coated coverslips. The peak intensity (I_max) was determined by finding a maximum value using Excel software (Microsoft). To determine the extent of calcium response, we calculated the decrease in intensity after the maximal response. The early phase of the measurement (an increase in intensity) represents a combination of the rise of background fluorescence due to cell spreading on the optical surface and of the specific sensor fluorescence caused by increased calcium in the cytosol. The kinetics of fluorescence decay was calculated as a ratio I_max/I_max+5min and I_max/I_max+10min. Higher values indicate a more rapid decay of the fluorescence, which indicates stronger response. Values of I_max+5min and I_max+10min represent the fluorescence intensity 5 and 10 minutes, respectively, after the maximum calcium response (I_max) was reached. To avoid the impact of the intensity fluctuations, we calculated I_max), I_max+5min and I_max+10min as an average of 10 images (frames).

### Characterization of T cell surface in contact with a coated coverslip and cell viability measurements

#### Microscope setup

For imaging of living T cells forming contact with a coated coverslip and for determining the viability of T cells immobilized on a coated coverslip, a home-build inverted microscope system (IX71, Olympus) equipped with 150 mW 488 nm and 150 mW 561 nm lasers (Sapphire, Coherent) was used. Fluorescence emission for cell surface-contact analysis and viability assay was detected by an EMCCD (iXon DU-897, Andor) and a sCMOS (Zyla-4.2-CL10, Andor) cameras, respectively. Two acousto-optic tunable filters (AOTFnC-400.650-TN, AA Optoelectronics) provided fast switching and synchronization of lasers with a camera. For TIRF imaging, the 488 nm laser beam was focused onto the back focal plane of an objective (UApoN 100x, NA = 1.49, Olympus). TIRF illumination was achieved using a manual micrometer-scale tilting mirror mechanism in the excitation pathway (Thorlabs). For cell viability measurements, the laser light was defocused to achieve a homogenous illumination at the objective sample plane (UPlanSApo 10x, NA = 0.4, Olympus). The system was controlled using the μManager software (version 1.4.22; ref.^93^).

#### Analysis of the T cell surface in contact with a coated coverslip

For T cell-coverslip contact analysis, Jurkat T CD4-KO cells were transfected with pXJ41-CD4-wt-eGFP plasmid 24-hours prior to the measurement. Cells were then harvested and transferred into color-free RMPI medium (Gibco) supplemented with 2 mM L-glutamine (Lonza) pre-warmed to 37°C. In parallel, cleaned coverslips were mounted for imaging into a ChamLide holder equipped with tubing adaptors (Live Cell Instruments), coated as described above and washed with ultrapure water. Immediately after coating fluid removal through the attached tubing, cells were injected into the ChamLide chamber attached to the microscope stage. Landing cells were selected under transmitted light and the fluorescence was recorded on EMCCD camera using TIRF illumination for 20 minutes. Images were acquired in one second intervals with 50 ms exposure time. The camera EM gain was set to 100. The experiment was performed at 37°C using the environmental chamber (OKO lab) with controlled temperature. Acquired images were processed using standard functions (ROI Manager, Plot Profile) of Fiji/ImageJ software (version 1.52p; ref.^79^).

#### T cell viability assay

To evaluate the viability of cells in contact with coated coverslips, Jurkat T cells were transfected with mTurquoise-Farnesyl-5 plasmid (#55551, Addgene) 24 hours prior to the assay. This was done to mimic the conditions used for all other experiments in this work but to avoid interference of the protein fluorescence with the dyes used for the assay. For the measurement, cells were washed with PBS, transferred into color-free RPMI (Gibco) medium supplemented with 2 mM *L*-glutamine (Lonza), 10% fetal bovine serum (Gibco), 10 mM HEPES and Non-essential amino-acid mixture (Gibco) containing 25 μg/ml (w/v) calcein-AM dye and incubated for 10 minutes at 37°C in the CO_2_ incubator. Cells were then transferred into a fresh, color-free RPMI medium supplemented with 2 mM *L*-glutamine and 7AAD (7-Amino-Actinomycine D; 1:20 dilution) viability stain (eBioscience) and incubated for another 5 minutes at 37°C in the CO_2_ incubator. After loading with the dyes, cells were injected onto the coated coverslips in ChamLide chamber using attached tubing. Imaging was performed for 30 minutes (with 10-minute acquisition intervals) from the time of the fist cell-coverslip contact detection. Two randomly selected ROIs were selected for each time point to avoid the effect of the fluorophore photo-destruction. Calcein and 7AAD dyes were excited separately by 488 nm and 561 nm lasers, respectively, using defocused light to achieve homogenous illumination. Cells were also imaged under transmitted light to control their morphology. To avoid the impact of fluorescence fluctuations, five frames were collected for each channel, ROI and time point. The frames were then summed to generate a collate image for further quantitative analysis. Data analysis was performed in Fiji/ImageJ (version 1.52p, ref.^79^). Cells were counted by determining local maxima for each channel. The final result was calculated as a percentage of 7AAD positive (dying) cells in all detected cells (calcein positive). The experiment was performed at 37°C using the environmental chamber (OKO lab) with controlled temperature.

### Atomic force microscopy

Atomic Force Microscopy (AFM) topography images were collected with Dimension Icon AFM (Bruker Instruments). All images were measured in the Peak Force Tapping mode for fluids, using a probe holder for fluid operation and Scanasyst-Fluid probes (Bruker) with a tip radius of 20 nm and a spring constant of 0.7 N/m.

12 mm diameter coverslips were coated with PLL and glycine as described above. A bare coverslip was prepared as a reference. The AFM probe was submerged in the solution in order to land on the coverslip surface. Due to a delicate nature of the samples and to avoid long measurement times, setpoint and number of lines were set to 400 pN and 256 × 256 lines respectively, with a scan rate of 1 Hz. Images with scan sizes of 1, 2 and 10 μm were collected.

## Acknowledgements

We would like to thank Peter Kapusta, Silke Kerruth and Harsha Mavila for technical assistance and professional advice. We are grateful to Markus Sauer (University of Würzburg) for support of the project. C.F. would like to thank Laure Plantard and Jan Peychl from the MPI-CBG LMF for technical support. M.C. acknowledges funding from Czech Science Foundation (19-0704S), S.vdL. acknowledges funding from Academy of Medical Sciences/the British Heart Foundation/the Government Department of Business, Energy and Industrial Strategy/the Wellcome Trust Springboard Award (SBF003\1163). The measurements at the Imaging Methods Core Facility in BIOCEV, Vestec, Czech Republic were supported by MEYS CR grant Z.02.1.01/0.0/0.0/16_013/0001775.

## Contributions

C.F., S.vdL. and M.C. conceived the study. C.F., T.C., Z.K., D.G. and D.A.H. performed the experiments, C.F., T.C., D.G., S.vdL. and M.C. analyzed the data. A.R. measured and analyzed the AFM data. O.F. and T.B. provided research advice. C.F., S.vdL. and M.C. wrote the paper. All authors reviewed and approved the manuscript.

## Additional information

Supplementary Information accompanies this paper at …

## Supplementary Information

**Supplementary Figure S1.**
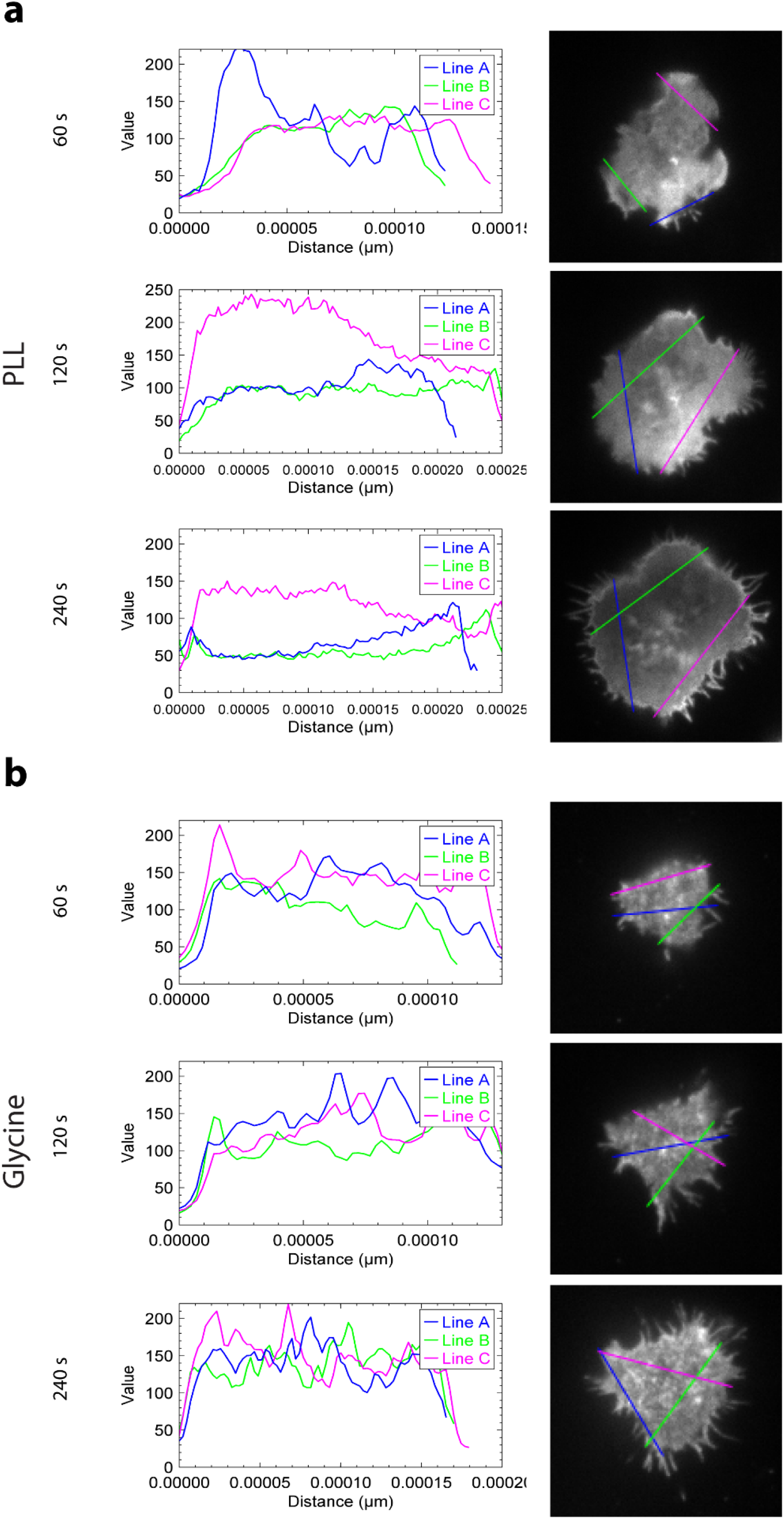
The impact of coverslip coating on cell surface morphology. Two-dimensional images of T cell-coverslip contact sections with indicated line selections (right panels) and the corresponding intensity line profiles (left panels). Cells were transfected with CD4-GFP, transferred to the chambers with PLL-(**a**) or glycine-coated (**b**) coverslips and living cells were imaged using TIRF microscopy at 1 fps. Snapshots acquired 60, 120 and 240 s after the first contact of the cell with the coverslip are presented.

**Supplementary Figure S2.**
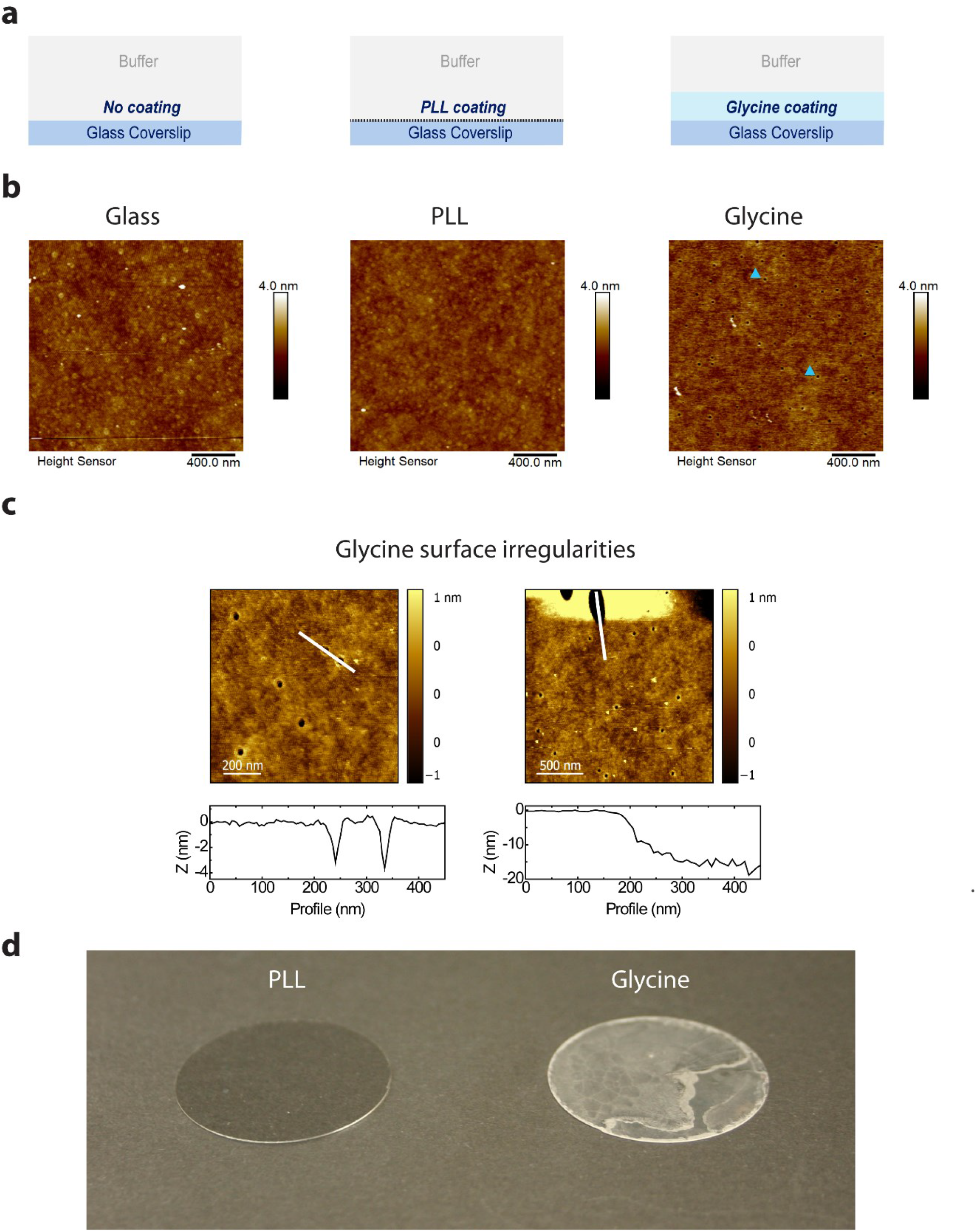
Glycine forms a narrow, gel-like structure on glass coverslips. **a)** Schematic illustration of the coverslip coatings tested using atomic force microscopy (AFM). **b)** Comparison of the AFM topography images of the coverslip surface without further coating (left panel) and after coating with poly-*L*-Lysine (PLL; middle panel) or glycine (right panel). Blue arrowheads in the right-hand panel indicate two examples of small topography features (holes), which populate the surface of glycine-coated coverslip. **c)** Magnified AFM topography images on a glycine-coated coverslip (upper panels) with respective height profiles (lower panels) taken in the area with the smaller, almost perfectly circular holes (diameter ~30 nm, depth ~3–4 nm on the average; left panels), and in the area with a larger hole with the diameter >200 nm and depth >20 nm (right panels). **d)** Glycine coating of glass coverslips (right) forms a white precipitate after a brief drying, indicating a hydrogel formation. No such precipitate can be observed on the PLL-coated coverslips (left), which do not differ from the uncoated coverslips (not shown).

**Supplementary Figure S3.**
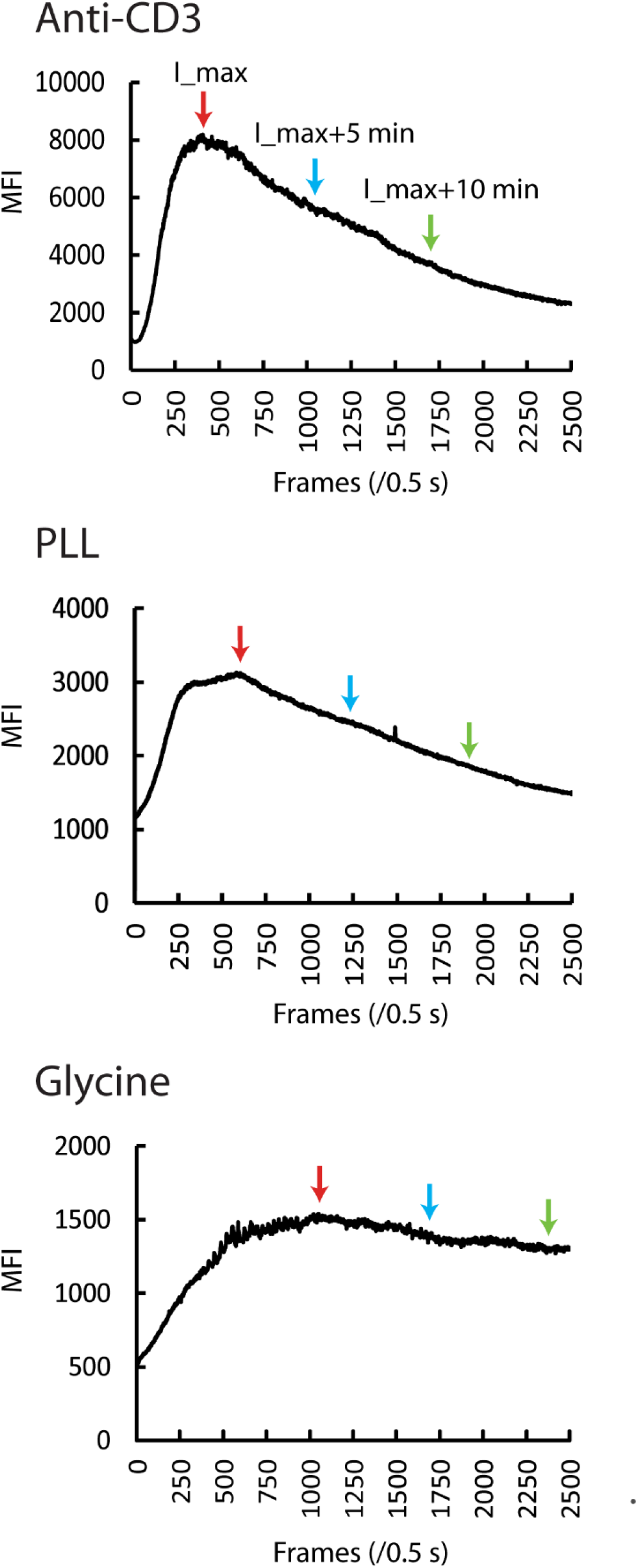
Calcium response measurements on coverslips functionalized to stimulate (OKT3 antibody) or immobilize T cells (PLL and glycine). Representative time profiles of mean fluorescence intensity measured in T cells transfected with calcium sensor (GCaMP6f_u_) using TIRF microscopy (2 fps) and illumination with 488 nm laser line. 11 cells were imaged on OKT3-antibody (upper panel), 16 cells on PLL-(middle panel) and 12 cells on glycine-coated (lower panel) coverslips. Cell footprints were manually selected at the time of maximum signal detection. Red arrows indicate maximum mean intensity (I_max), blue the signal detected 5 minutes later (I_max+5 min) and green the signal detected after another 5 minutes (I_max+10 min). The calcium response of cells interacting with the functionalized surface was calculated as I_max/I_max+5min and I_max/I_max+10min as summarized in Fig.1e.

**Supplementary Figure S4.**
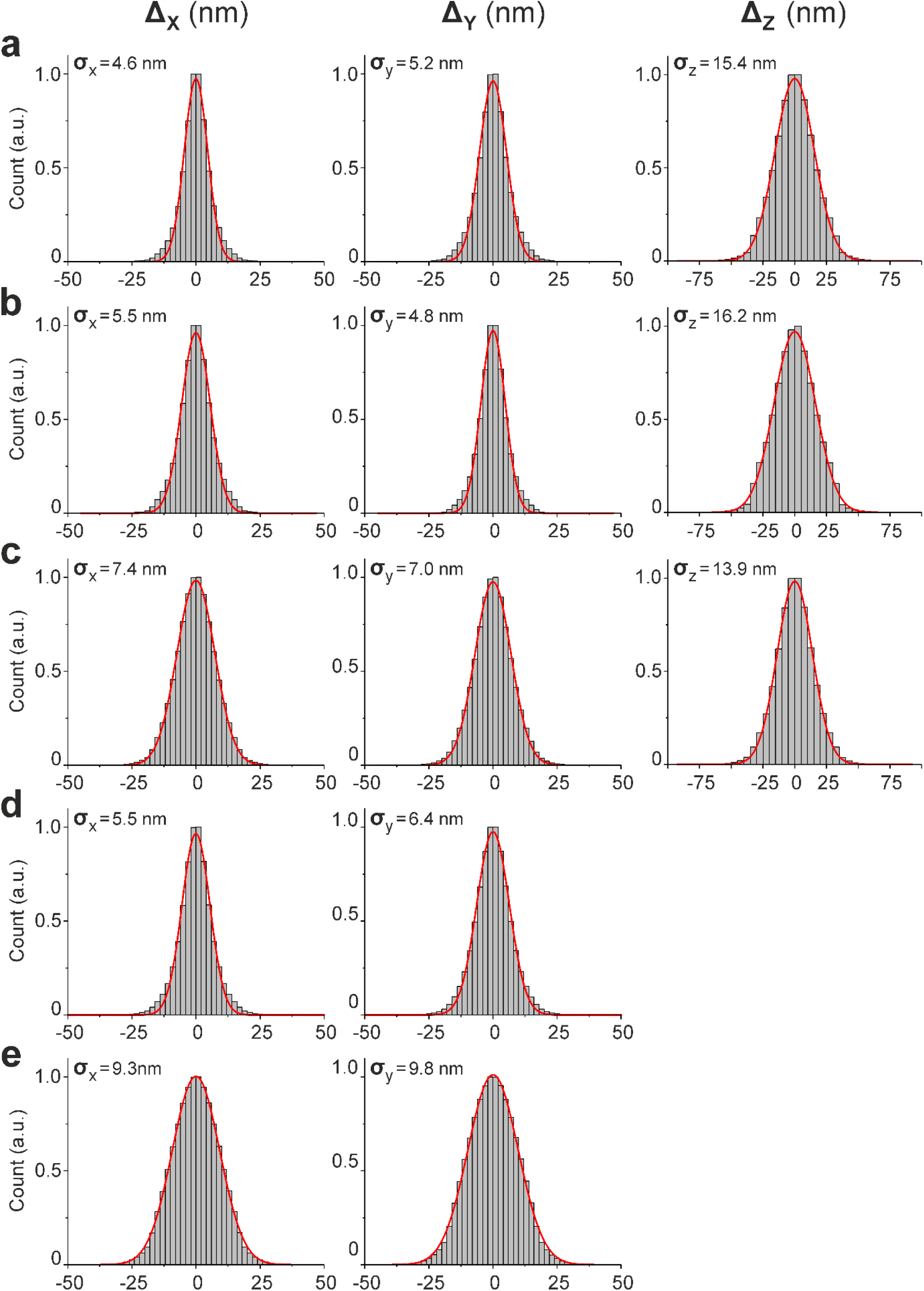
Localization precision. Normalized histograms of spatio-temporal nearest neighbour tracks, constituting the three-dimensional localization precision of the dTRABI and two-color SMLM measurements (see **Methods**). x-, y- and z- were individually determined and are stated as the standard deviation of the respective distribution as numerical value σ. Sample and imaging modalities: **a)** dTRABI of CD4 WT (Alexa Fluor 647) **b)** dTRABI of CD4 CS1 (Alexa Fluor 647) **c)** dTRABI of CD45 (Alexa Fluor 647) **d)** two-dimensional *d*STORM of CD45 (Alexa Fluor 647) **e)** two-dimensional PALM of CD45 (mEos2).

**Supplementary Figure S5.**
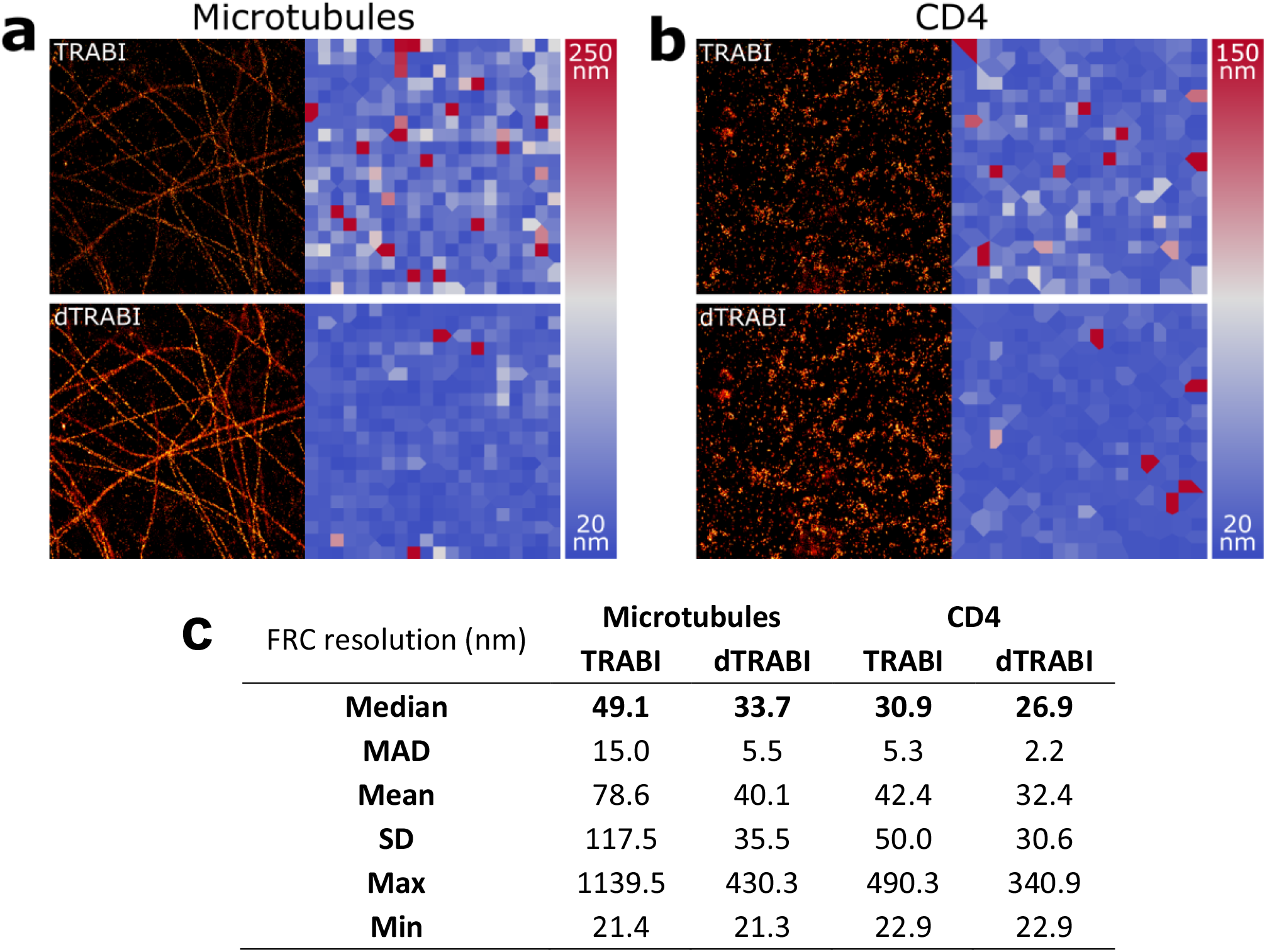
Resolution improvement of dTRABI compared to TRABI determined by Fourier Ring Correlation (FRC). **a**) Representative three-dimensional dSTORM images of microtubules (data from ref.^1^). The three-dimensional localization sets were derived according to either the standard TRABI-Biplane algorithm (TRABI, *top*) or the new dTRABI approach (*bottom*). The according FRC maps, visualizing the local FRC-resolution, are depicted on the right. We selected the median FRC resolution of the images as the most robust resolution metric, which was derived to 49 nm (TRABI) and 34 nm (dTRABI). dTRABi improves the structural resolution in a medium density sample by 30 percent. **b**) Representative three-dimensional dSTORM images of CD4 (data reanalyzed from **Fig. 5a**). The three-dimensional localization sets were derived according to either the standard TRABI-Biplane algorithm (TRABI, *top*) or the dTRABI approach (*bottom*). The according FRC maps are depicted on the right. The median FRC resolution of the images was derived to 31 nm (TRABI) and 27 nm (dTRABI). dTRABI improves the structural resolution in a low-density sample by 13 percent. FRC analysis was performed with NanoJ-Squirrel^2^, additional relevant FRC metrics are listed in **c**). Scale bars, 1 μm.

**Supplementary Figure S6.**
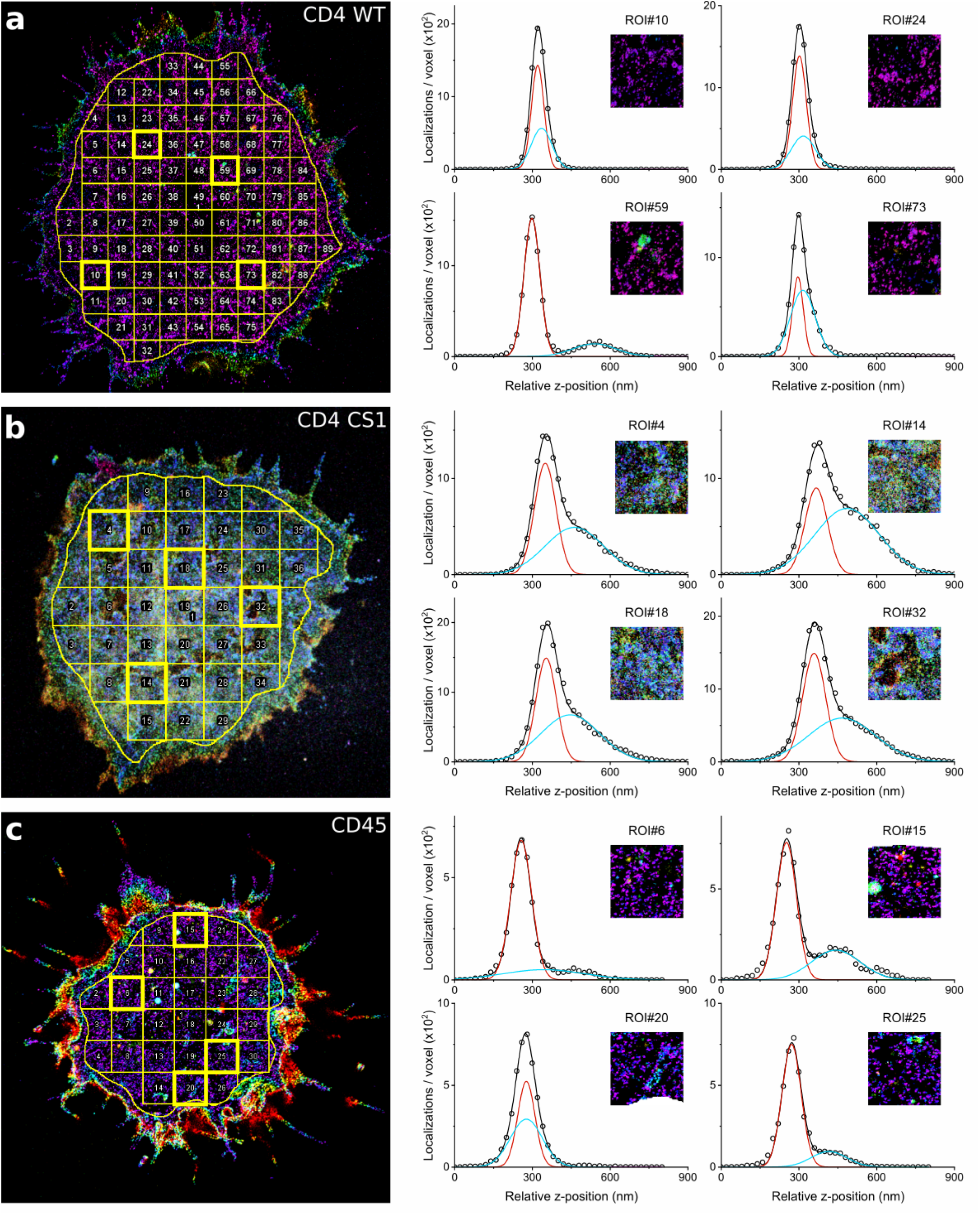
Segmentation of cells. For quantitative axial analysis of surface receptors, the interior of each cell footprint was first manually selected (ROI#1) to avoid the impact of cell edges. Afterwards, the area was further segmented into squared 2 μm × 2 μm ROIs. In border areas, ROIs were kept if the area was ≥75% of 4 μm^2^ (see **Methods**). **a**) Segmentation and quantitative axial analysis for CD4 WT. *Left*: The cell as shown in Fig. 5a segmented, *right*: example ROIs and Gaussian fitting for quantitative analysis of receptor z-distribution as in Fig.5e-g. **b**) Segmentation and quantitative axial analysis for CD4 CS1. *Left*: The cell as shown in Fig. 5c segmented, *right*: example ROIs and Gaussian fitting for quantitative analysis of receptor z-distribution. **c**) Segmentation and quantitative axial analysis for CD45. *Left*: The cell as shown in Fig. 4b segmented, *right*: example ROIs and Gaussian fitting for quantitative analysis of receptor z-distribution. Selected ROIs are depicted with a bold frame in the whole-cell images on the left-hand side. In graphs with the axial receptor distribution, black circles represent raw data, black line the bi-Gaussian fit, which is the sum of two Gaussians as depicted in red and blue.

**Supplementary Figure S7.**
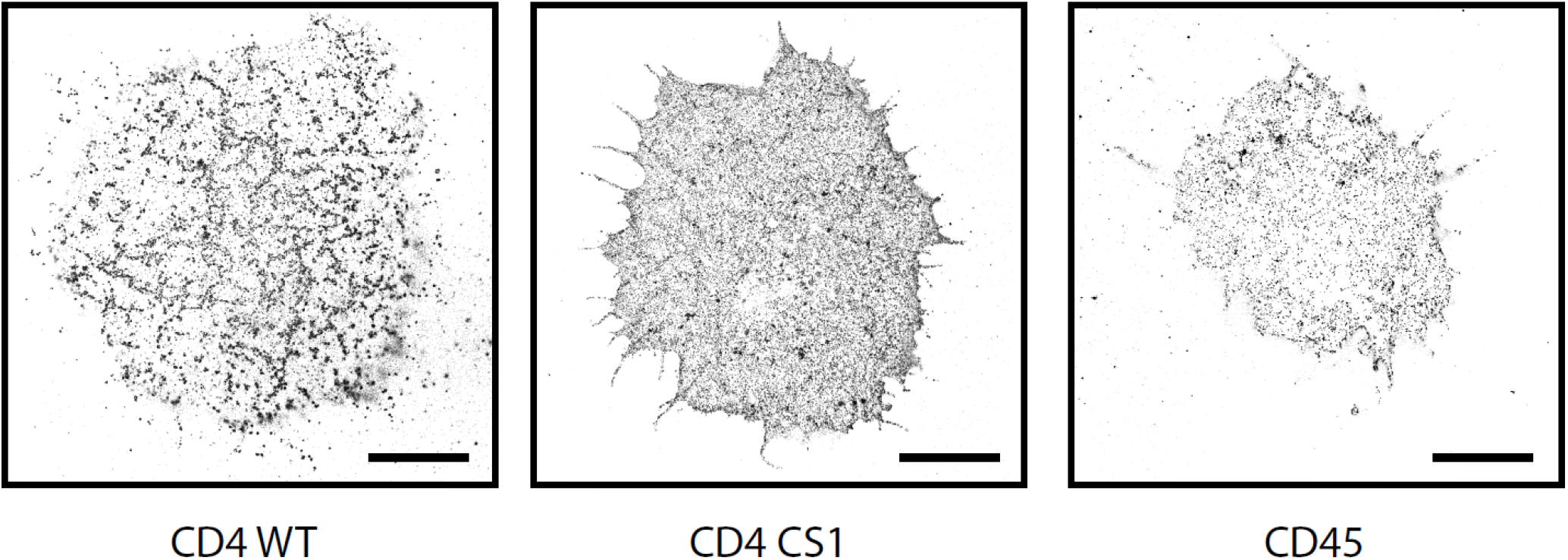
Two-dimensional SMLM of tested T cell receptors immobilized on glycine-coated coverslips. dSTORM images of CD4 WT (left), CD4 CS1 (middle) and CD45 (right) on the surface of unstimulated T cells. The two-dimensional images indicate different nanoscopic organization of the receptors but cannot be analyzed quantitatively due to a three-dimensional character of the T cell surface. Scale bars, 5 μm.

**Supplementary Figure S8.**
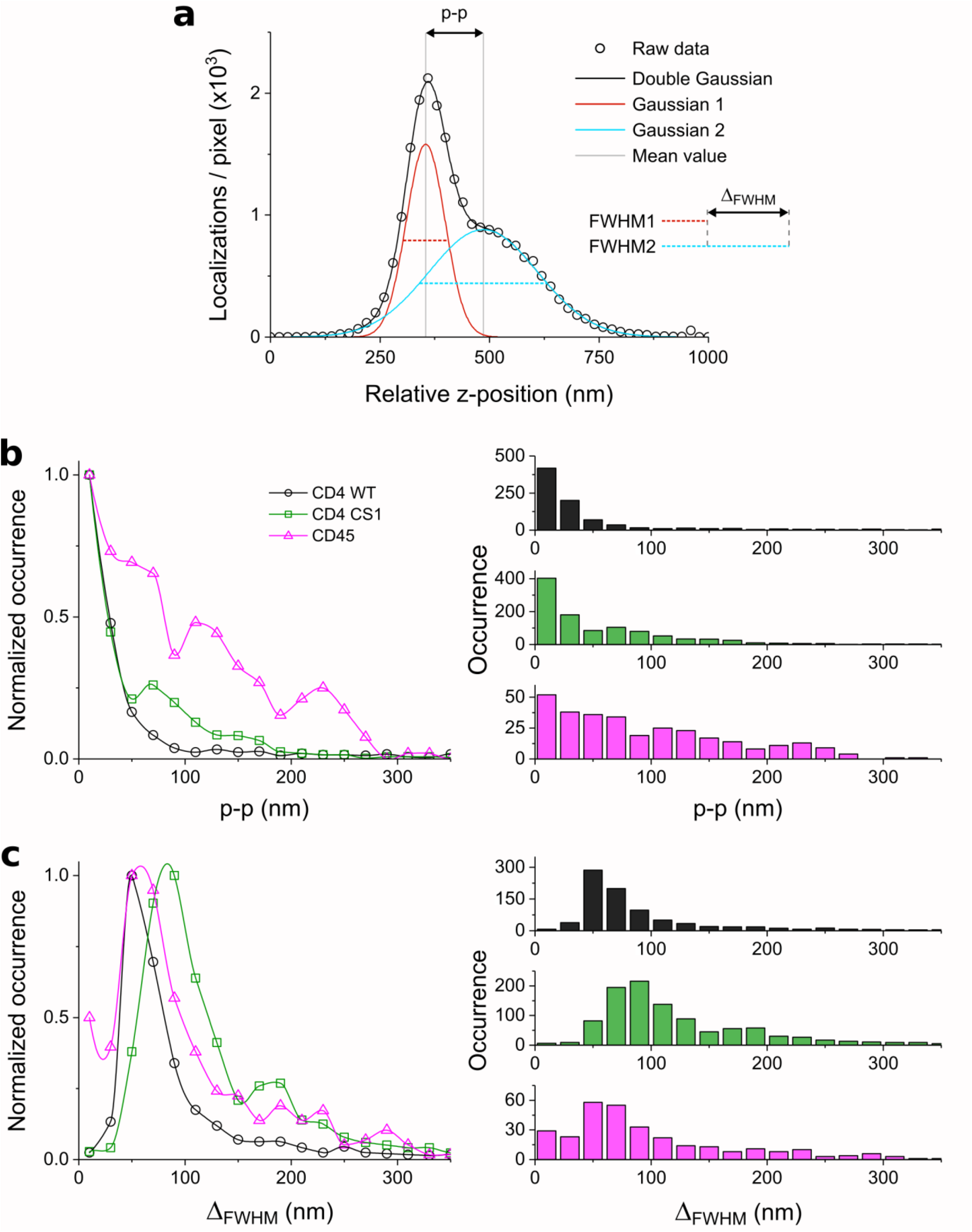
Quantitative analysis of axial receptor distribution. **a**) The principle of data analysis using a bi-Gaussian fit to derive the quantitative parameters peak-to-peak distance (p-p) and width difference (Δ_FWHM_). **b**) p-p, which represents the absolute value of the difference the mean values as indicated in a) (p-p = mean2-mean1). **c**) Δ_FWHM_, which represents the absolute value of the difference between the FWHM values of the two Gaussians as indicated in a) (Δ_FWHM_ = FWHM2-FWHM1). The normalized distributions are plotted on the left and histograms with absolute values on the right-hand side. In b and c, black represents data for CD4 WT, green for CD4 CS and magenta for CD45. All data points were spline interpolated to guide the eye. For CD4 WT 21 cells, CD4 CS1 18 cells and CD45 13 cells were analyzed.

**Supplementary Figure S9.**
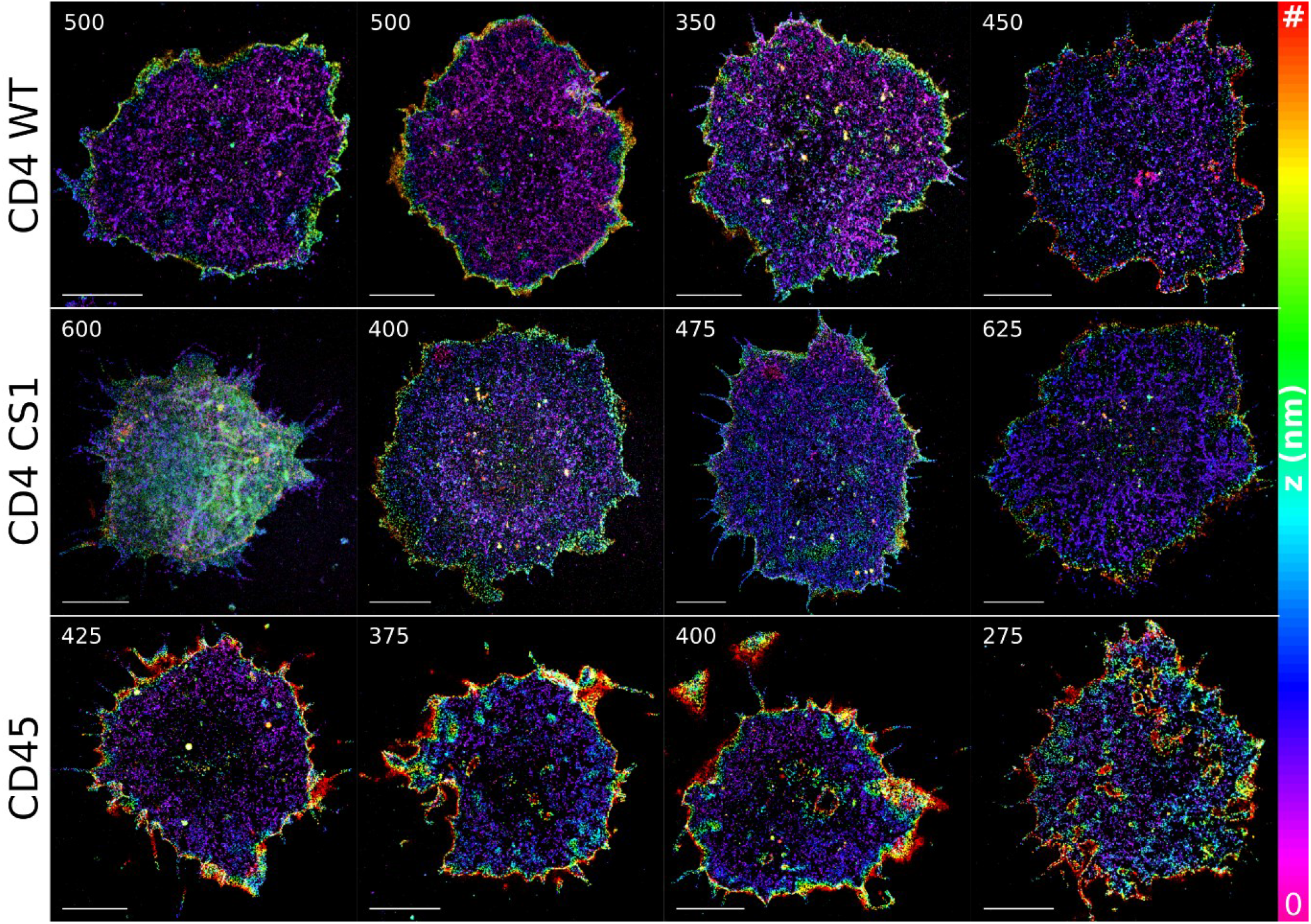
Exemplary selection of dTRABI images for CD4 WT, CD4 CS1 and CD45. The maximum z-value is depicted for each cell (top left corner), e.g. for the cell in the upper right corner the z-range is from 0 to 450 nm. Marker ‘#’ in the color-bar means the maximum z-value for each cell. In total, 21, 18 and 13 cells were analyzed for CD4 WT, CD4 CS1 and CD45, respectively. Scale bars, 5 μm.

**Supplementary Figure S10.**
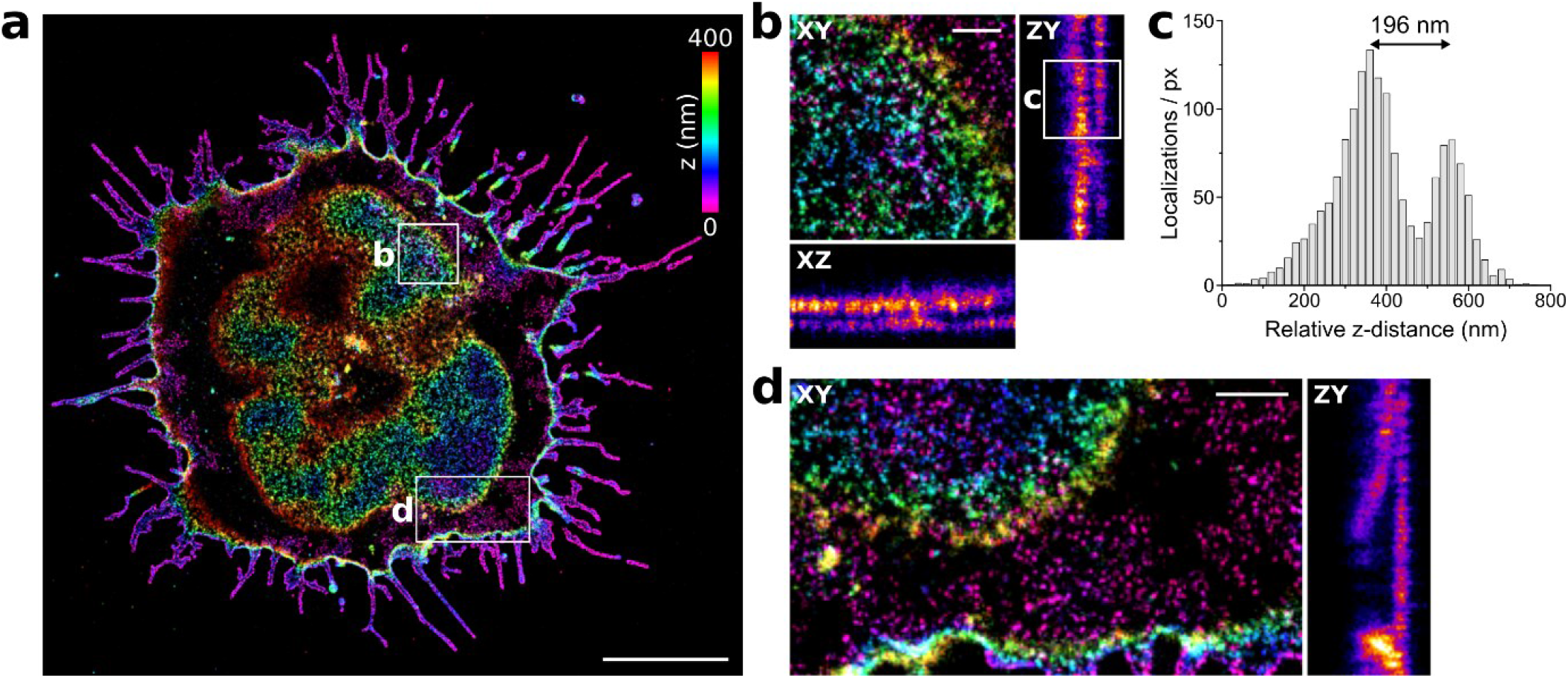
Cell surface receptor nanotopography on a dying cell with extensive three-dimensional deformations visualized by three-dimensional dTRABI imaging. **a**) Three-dimensional dTRABI image of CD4 with selected ROIs exhibiting broad z-distribution of receptor localizations. **b)** Magnified x-y, x-z and z-y projections of the ROI indicated in a). **c)** The axial distribution of localizations as depicted in the y-z plot in b). **d)** Magnified x-y and z-y projections of the ROI indicated in a). Scale bars, a) 5 μm (x, y), b, d) 500 nm (x, y, z).

**Supplementary Figure S11.**
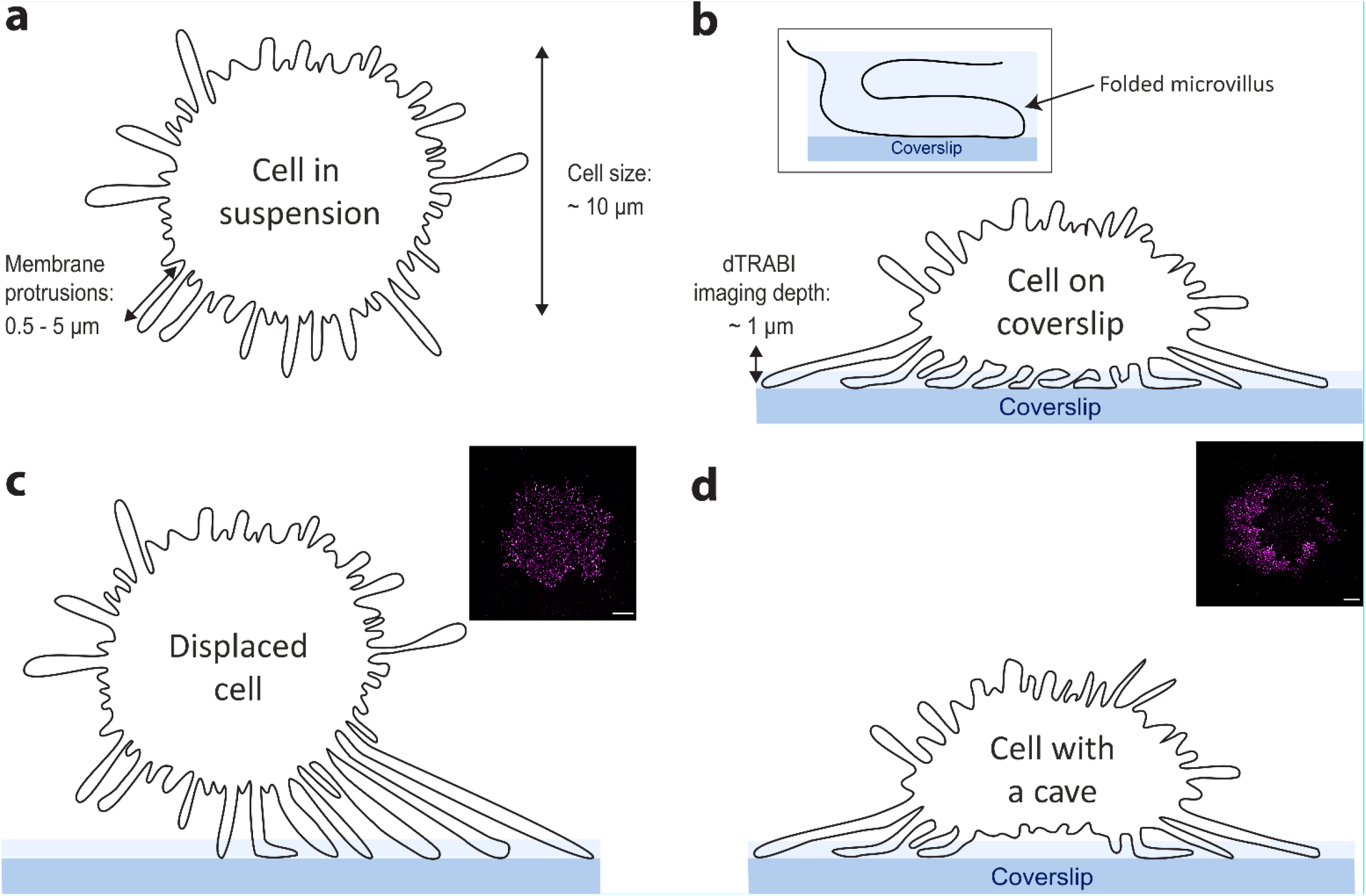
T-cell surface morphology on glycine-coated coverslips. **a)** Schematic illustration of a T cell in suspension. The cell is covered with numerous membrane protrusions of different sizes. The size of Jurkat T cells is, on average, 10 μm in diameter, the protrusions, even though a majority are small (0.5–2 μm length), can extend up to 5 μm from the cell body. **b)** After landing on a glycine-coated coverslip, membrane protrusions of T cells fold under the cell body but are not rapidly removed as on PLL. **c-d)** With the axial penetration depth of dTRABI being ~1 μm, we were unable to detect receptors at the plasma membrane basis of T cells exhibiting displacement of the cell body from the contact site (c) or those with the cavity formed between the cell body and the coated coverslip. The footprints of such cells show discontinuous labeling or a hole in the middle of the image (see the inserts). The blue stripes represent glass coverslip, the light blue stripes above, the glycine layer (not to scale).

## Supplementary Movies

**Supplementary Movie 1 Live cell TIRF microscopy of CD4-GFP in T cells landing on PLL**-coated measured at 37°C. Representative cell as in Fig. 1c shown. Acquisition rate: 1 fps; video fame rate: 15 fps; scale bar: 5 μm. The image sequence was corrected for the photobleaching; the brightness and contrast were leveled using plugins of Fiji software.

**Supplementary Movie 2. Live cell TIRF microscopy of CD4-GFP in T cells landing on glycine**-coated coverslips measured at 37°C. Representative cell as in Fig. 1d shown. Acquisition rate: 1 fps; video fame rate: 15 fps; scale bar: 5 μm. The image sequence was corrected for the photobleaching; the brightness and contrast were leveled using plugins of ImageJ/Fiji software.

